# N-MYC regulates Cell Survival via eIF4G1 in inv(16) Acute Myeloid Leukemia

**DOI:** 10.1101/2023.03.03.531018

**Authors:** Philomina Sona Peramangalam, Sridevi Surapally, Shikan Zheng, Robert Burns, Nan Zhu, Sridhar Rao, Carsten Muller-Tidow, John A. Pulikkan

## Abstract

c-MYC and N-MYC are critical regulators of hematopoietic stem cell activity. While the role of c-MYC deregulation is studied in detail in hematological malignancies, the importance of N-MYC deregulation in leukemogenesis remains elusive. Here we demonstrate that N-MYC is overexpressed in acute myeloid leukemia (AML) cells with chromosome inversion inv(16) and crucial to the survival and maintenance of inv(16) leukemia. We identified a novel *MYCN* enhancer, active in multiple AML subtypes, essential for MYCN mRNA levels and survival in inv(16) AML cells. We also identified eukaryotic translation initiation factor 4 gamma 1 (eIF4G1) as a key N-MYC target that sustains leukemic survival in inv(16) AML cells. Ours is the first report to demonstrate the oncogenic role of eIF4G1 in AML. Our results reveal a mechanism whereby N-MYC drives a leukemic transcriptional program and provide a rationale for the therapeutic targeting of the N-MYC/eIF4G1 axis in myeloid leukemia.

## Introduction

The core binding factor (CBF) is a heterodimeric transcription factor complex composed of DNA-binding RUNX proteins (encoded by one of three genes -RUNX1, RUNX2, and RUNX3) and the non-DNA-binding CBFβ protein. RUNX1 and CBFβ play essential roles in an overlapping and differentiation stage-specific manner in embryonic and adult hematopoiesis.^1,2^ The inv(16)(p13q22) result in the formation of a chimeric gene consisting of the 5’ portions of *CBFB* fused to the 3’ portion of the smooth muscle myosin heavy chain gene, *MYH11*, which encodes CBFβ–SMMHC fusion protein. Inv(16) is reported in 5-8 % of AML patients. CBFβ–SMMHC has a higher affinity for RUNX1 than the native CBFβ ^3,4^. CBFβ–SMMHC expressing embryos die at mid-gestation due to block in definitive hematopoiesis ^5^, which is very similar to the phenotype from Runx1 and Cbfb knockout embryos, ^1,2,^ suggesting CBFβ–SMMHC fusion protein is a dominant repressor of RUNX1. Studies using conditional knock-in mouse models demonstrated that CBFβ–SMMHC deregulates hematopoietic stem cell (HSC) differentiation in adult hematopoiesis by expanding short-term HSCs and myeloid progenitor cells. ^6-8^ These aberrant myeloid progenitors with pre-leukemic potential upon acquisition of secondary cooperating mutations induce AML. ^9-11^ Recent attempts to inhibit protein-protein interaction between RUNX1 and CBFβ–SMMHC via small molecule inhibitor AI-10-49 demonstrated promising results in selectively inducing apoptosis in primary human inv(16) AML cells and a genetic mouse model for inv(16). ^12,13^ Further, we demonstrated that AI-10-49 induced cell death is partly mediated by repression of c-MYC expression in inv(16) AML cells. ^14^

The MYC family genes include three closely related genes *MYC* (encoding c-MYC), *MYCN* (encoding N-MYC), and *MYCL* (encoding L-MYC). ^15^ While c-MYC function is widely investigated in hematological malignancies, the function of other MYC family members in inv(16) AML, and AML in general, remains poorly understood. Here we demonstrate that AI-10-49 treatment in inv(16) AML cells downregulated MYCN transcript levels. Utilizing primary human inv(16) AML cells and patient-derived xenograft (PDX) model, we demonstrate that N-MYC is required for the maintenance of inv(16) AML survival. Furthermore, by comparing N-MYC transcriptional targets with gene expression changes in human inv(16) AML, we identified eukaryotic translation initiation factor 4 gamma 1 (eIF4G1) as a key N-MYC target in leukemic survival. Collectively, our results illustrate that the oncogenic N-MYC/eIF4G1 axis is instrumental in inv(16) AML leukemogenesis.

## Results

### MYCN is upregulated in inv(16) AML

AI-10-49 treatment results in transcriptional changes, including c-MYC repression, and triggers apoptosis in inv(16) AML cells. ^14^ To understand the potential role of other MYC family members, we re-analyzed previous RNA-seq data. ^14^ We found that MYCN transcript is downregulated with AI-10-49 treatment in inv(16) AML cell line ME-1 (**Figure 1A-B and Supplemental Figure 1A**). MYCL transcript is not expressed in ME-1 cells (data not shown). Similar to the repression of MYCN at transcript levels, we also observed the downregulation of N-MYC protein levels with AI-10-49 in inv(16) AML cells (**Figure 1C and Supplemental Figure 1B**). Downregulation of MYCN expression by AI-10-49 suggested that MYCN overexpression could be a critical step in inv(16) leukemogenesis. To test this, we analyzed MYCN transcript levels in human AML patient samples. Our data demonstrated that MYCN transcript levels were upregulated in primary inv(16) AML patient samples compared to healthy control samples (**Figure 1D-E**). Bromodomain inhibitors have been reported to repress MYC and MYCN in cancer cells. ^16,17^ We have previously shown that bromodomain inhibitor JQ1 downregulates MYC transcript and synergizes with AI-10-49 in inducing apoptosis in inv(16) leukemic cells. We found that JQ1 treatment in inv(16) AML cells induces repression of MYCN transcription (**Supplemental Figure 1C**). This suggests that in addition to c-MYC, N-MYC may have important roles in apoptosis induction by JQ1 in inv(16) AML cells in the previous study. ^14^

**Figure 1.**
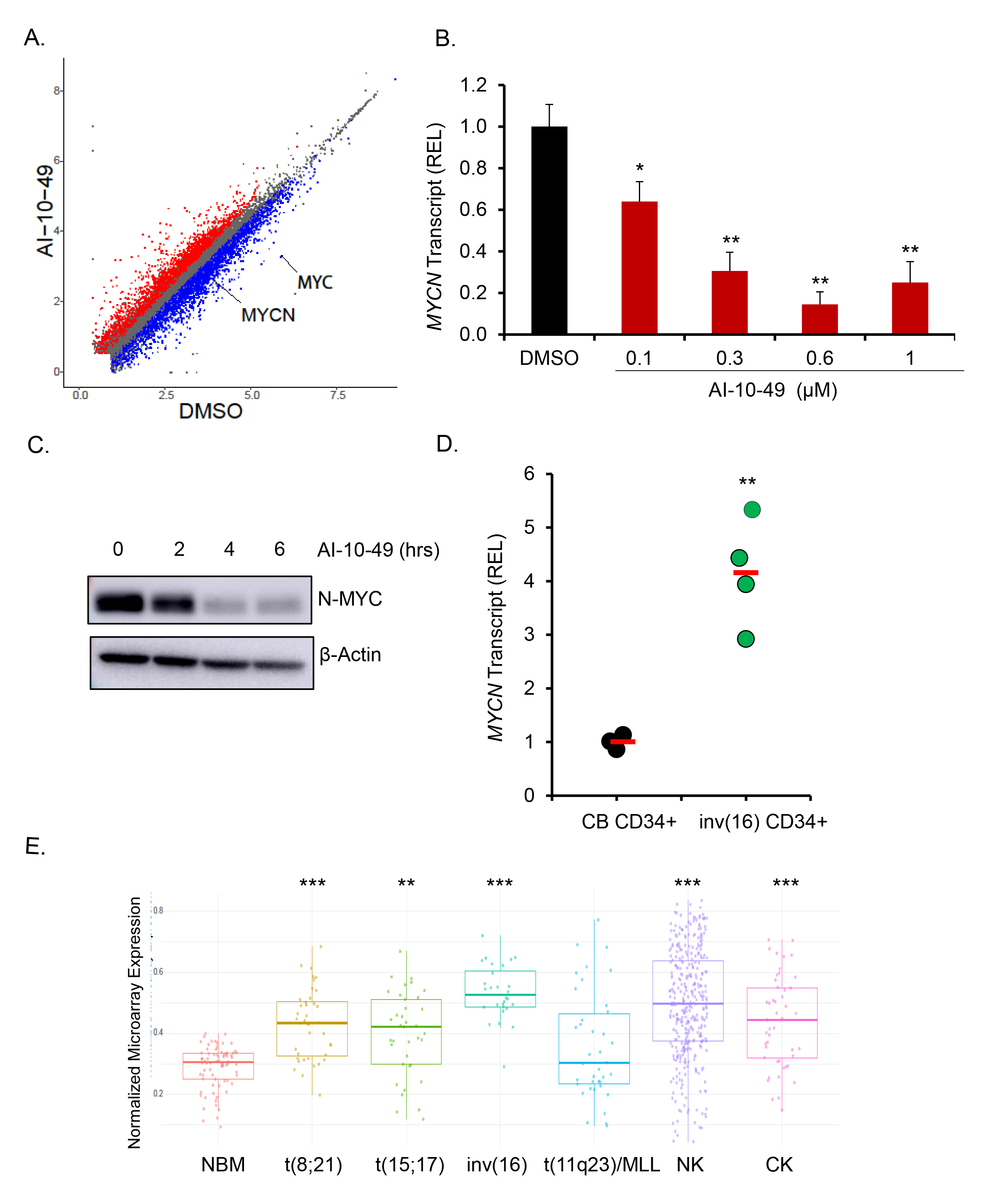
MYCN is upregulated in inv(16) AML. (**A**). Scatter plot of differentially expressed genes in RNA-seq analysis between DMSO- and AI-10-49-treated ME-1 cells (>2-fold change; FDR < 0.01). Genes significantly changed are colored in red and blue for upregulated and downregulated, respectively. c-MYC and N-MYC are highlighted. (**B**). N-MYC transcript levels in DMSO / AI-10-49 treated (6 hrs) ME-1 cells by Real Time RT-PCR. Histogram representative of triplicate experiments. (**C**). N-MYC protein levels in AI-10-49 treated (1 μM) ME-1 cells by western blot. (**D**). MYCN transcript levels in human cord blood CD34+ cells (CB CD34+) and primary human inv(16) AML CD34+ cells (inv(16) CD34+). Each symbol represents the average of a triplicate experiment from one sample, and the average value of the group is shown in red. (**E**). Normalized expression of MYCN transcript levels in Haferlach dataset extracted from Leukemia Gene Atlas (expression array) database. Error bars represent the SD. Significance was calculated using an unpaired t-test. *p < 0.05 or **p < 0.005 or ***p < 0.0005.

### N-MYC is required for inv(16) AML cell survival

To understand the functional role of N-MYC in inv(16) AML cells, we assessed whether the deletion of MYCN affects the survival of inv(16) AML cells. We applied CRISPR/Cas9 technology using a ribonucleoprotein (RNP) complex approach to delete *MYCN* in inv(16) AML cells. ME-1 cells transfected with Cas9 protein and three *MYCN* chemically modified guide RNAs (CM-gRNAs) produced efficient deletion of *MYCN* genomic regions (**Figure 2A and Supplemental Figure 2A-B**), resulting in a 95% reduction in N-MYC protein levels (**Figure 2B**). *MYCN* deletion in ME-1 cells significantly reduced cell viability due to apoptosis (**Figure 2C**). The onset of apoptosis was accompanied by cleavage of caspase 3 and poly ADP ribose polymerase (PARP) proteins in ME-1 cells (**Figure 2D**). Granulocyte differentiation markers such as CD11B, CD15, neutrophil elastase, and CSF1-R were not significantly changed during *MYCN* deletion (**Supplemental Figure 2C-F**), suggesting *MYCN* deletion induces apoptosis and not granulocytic differentiation in inv(16) AML cells. Taken together, these results indicate that inv(16) AML cells depend on N-MYC levels for survival.

**Figure 2.**
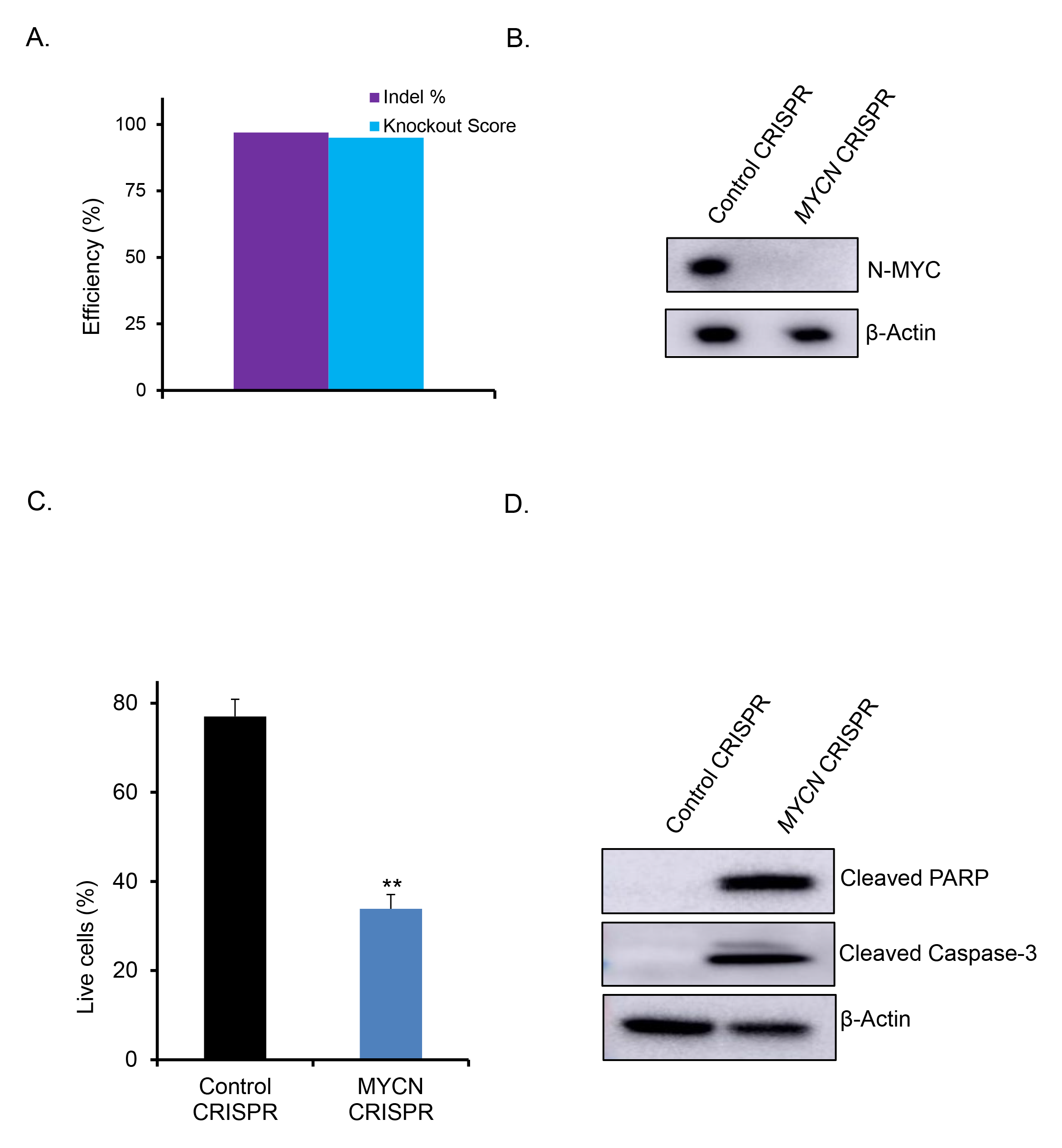
N-MYC is required for inv(16) AML cell survival. (**A**). ME-1 cells were transfected with Cas9 and control gRNA/ pool of 3 *MYCN* gRNAs [Synthego Gene Knockout kit V2] by RNP approach and analyzed editing efficiency by Inference of CRISPR editing (ICE). (**B**). N-MYC protein levels in control/ *MYCN* edited ME-1 cells by western blot. (**C**). Cell survival analysis in control/ *MYCN* edited ME-1 cells by Annexin V/7AAD assay. Histogram representative of triplicate experiments. (**D**). Cleaved PARP and cleaved Caspase-3 protein levels in control/ *MYCN* edited ME-1 cells by western blot. Error bars represent the SD. Significance was calculated using an unpaired t-test. **p < 0.005.

### N-MYC silencing effectively reduces AML burden and extends survival in the NSGS xenograft model

To further evaluate the role of N-MYC in inv(16) AML cell survival, we conducted *MYCN* genomic editing in primary human inv(16) AML cells. T cell-depleted primary inv(16) AML cells were transfected with Cas9 protein and three *MYCN* CM-gRNAs (**Figure 3A**), resulting in a 90% reduction in N-MYC protein levels (**Figure 3B**). *MYCN* deletion significantly reduced the clonogenic potential of primary inv(16) AML cells, as reflected by a marked decrease in colony number (**Figure 3C**). The requirement of N-MYC in inv(16) leukemia in vivo was tested by transplanting the edited leukemic cells into irradiated (280 cGy) NSGS [NOD/SCID-IL2RG– SGM3] mice ^18,19^ via tail vein injection. Engraftment efficiency of control and *MYCN* edited leukemic cells in the bone marrow five days after transplantation was similar between groups (**Supplemental Figure 3A-B**). AML burden estimated by the frequency of hCD45+ hCD33+ AML stem/progenitor cells was significantly reduced in the bone marrow and peripheral blood of mice transplanted with *MYCN* deleted inv(16) AML cells compared to control mice (**Figure 3D-F**), suggesting that N-MYC is required for the maintenance of inv(16) leukemic cells. In addition, the median leukemic latency in mice transplanted with *MYCN* deleted inv(16) AML cells was significantly extended from 70 days (control group) to 123 days (*MYCN* deleted group; p <0.005) (**Figure 3G**). Collectively, these results demonstrate that N-MYC is crucial for the survival and maintenance of inv(16) leukemia.

**Figure 3.**
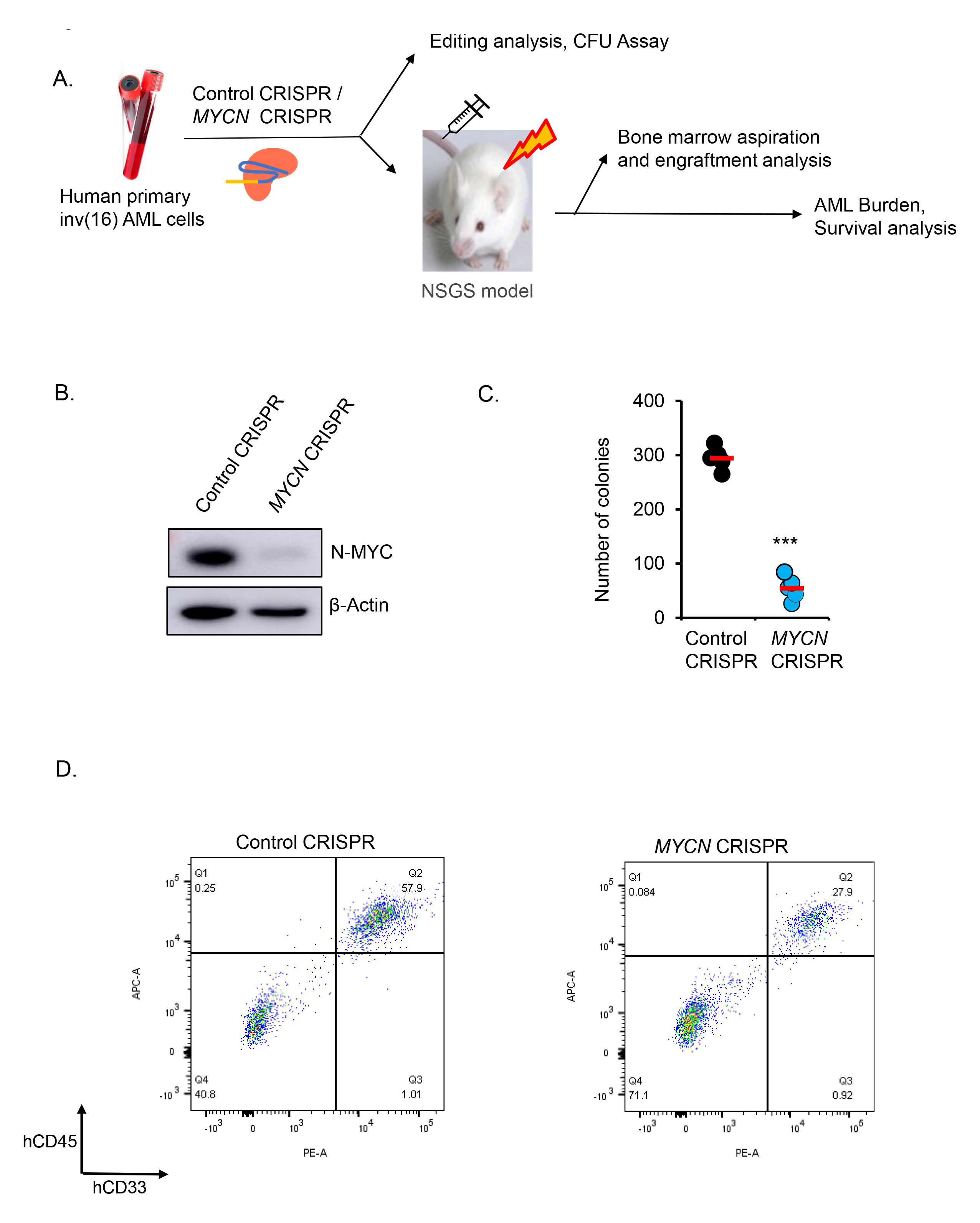
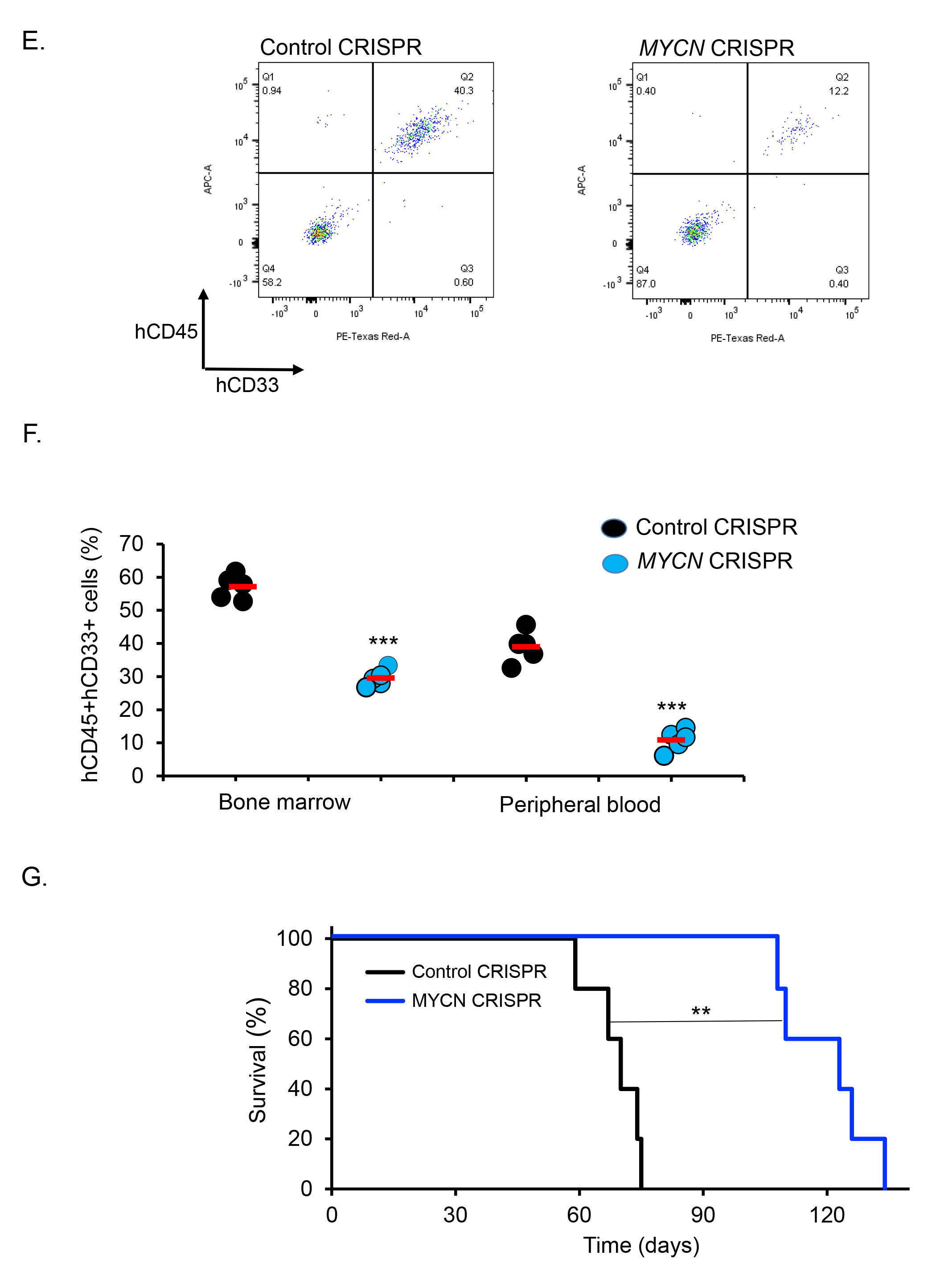
N-MYC silencing effectively reduces AML burden and extends survival in the NSGS xenograft model. (**A**). Schematic of *MYCN* editing by CRISPR/Cas9 RNP approach in primary human inv(16) AML cells and transplantation in NSGS mouse model. (**B**). N-MYC protein levels in control/ *MYCN* edited primary human inv(16) AML cells by western blot. (**C**). Colony counts for methylcellulose colony-forming assay performed upon 12 days of control/ *MYCN* edited primary inv(16) AML cells. Data representative of the four replicates. The average value of each group is shown in red. (**D-E**). Representative flow cytometry plots showing gating and frequency of hCD45+ hCD33+ cells in bone marrow (D) and peripheral blood (E) of NSGS mice transplanted with control/ *MYCN* edited primary inv(16) AML cells five weeks after transplantation. (**F**). Flow cytometric quantification of hCD45+ hCD33+ cells in NSGS mice transplanted with control/ *MYCN* edited primary inv(16) AML cells five weeks after transplantation. Each symbol represents a mouse. The average value of each group is shown in red. (**G**). Kaplan-Meier survival curve of NSGS mice transplanted with control/ *MYCN* edited primary inv(16) AML cells (n=5/group). Error bars represent the SD. Significance was calculated using an unpaired t-test (C, F) and log-rank test (G).**p < 0.005 or ***p < 0.0005.

### RUNX1 binds to an *MYCN* distal enhancer transcriptionally active in inv(16) AML cells

Distal enhancers regulate target gene transcription through chromatin loops that link enhancers with target gene promoters. ^20^ We have previously shown that AI-10-49 induces genome-wide RUNX1 occupancy in inv(16) AML cells. ^14^ Importantly, we have demonstrated that RUNX1 binding at *MYC* distal enhancers plays a critical role in c-MYC expression and survival in inv(16) AML cells. ^14^ We hypothesized that AI-10-49 mediated *MYCN* repression is due to direct RUNX1 binding at *MYCN cis*-regulatory elements, such as enhancers. To understand RUNX1 regulation of *MYCN*, we analyzed RUNX1 DNA binding profile and chromatin accessibility at the *MYCN* locus, previously reported in ME-1 cells. ^14^ Analysis of RUNX1 ChIP-seq peaks in AI-10-49 treated cells identified a genomic element located 26 kb downstream of *MYCN* transcription start site (TSS), named *RDME* (*<u>R</u>UNX1-<u>D</u>ependent <u>M</u>YCN <u>E</u>nhancer*) with increased RUNX1 binding (**Figure 4A**). Analysis of the genomic sequence of *RDME* identified a consensus RUNX binding site (TGYGGT). Previous studies have identified multiple enhancer elements (e1-e5) within 1.5 Mb downstream of *MYCN* TSS regulating MYCN expression in brain tumor cells. ^21^ Enhancer element e2 (*MYCN-e*2) contains a consensus RUNX binding site. We did not find any RUNX1 binding at *MYCN-e2* with AI-10-49 treatment in inv(16) AML cells (**Figure 4A**), suggesting it is irrelevant in hematopoietic cells. Enhanced RUNX1 binding at *RDME*, but not at *MYCN-e2* and the *MYCN* promoter, was validated by conducting ChIP-qPCR in ME-1 cells (**Figure 4B**) and primary human inv(16) AML cells (**Figure 4C**). H3K27ac, a histone mark representing transcriptionally active regions, was enriched at *RDME* and *MYCN* promoter (PR) and significantly reduced with AI-10-49 (**Figure 4A, D**), suggesting RUNX1 binding at *RDME* is resulting in transcriptional repression. Analysis of chromatin accessibility by ATAC-seq demonstrated that *RDME* is a highly accessible chromatin element (**Figure 4A**). Based on the ATCA-seq and H3K27ac ChIP-seq, *MYCN-e2* is transcriptionally inactive in inv(16) AML cells. H3 Lys 4 mono-methylation (H3K4me1) histone mark, which represents active enhancers, is enriched at *RDME,* suggesting *RDME* functions as a bonafide enhancer (**Figure 4E**).

**Figure 4.**
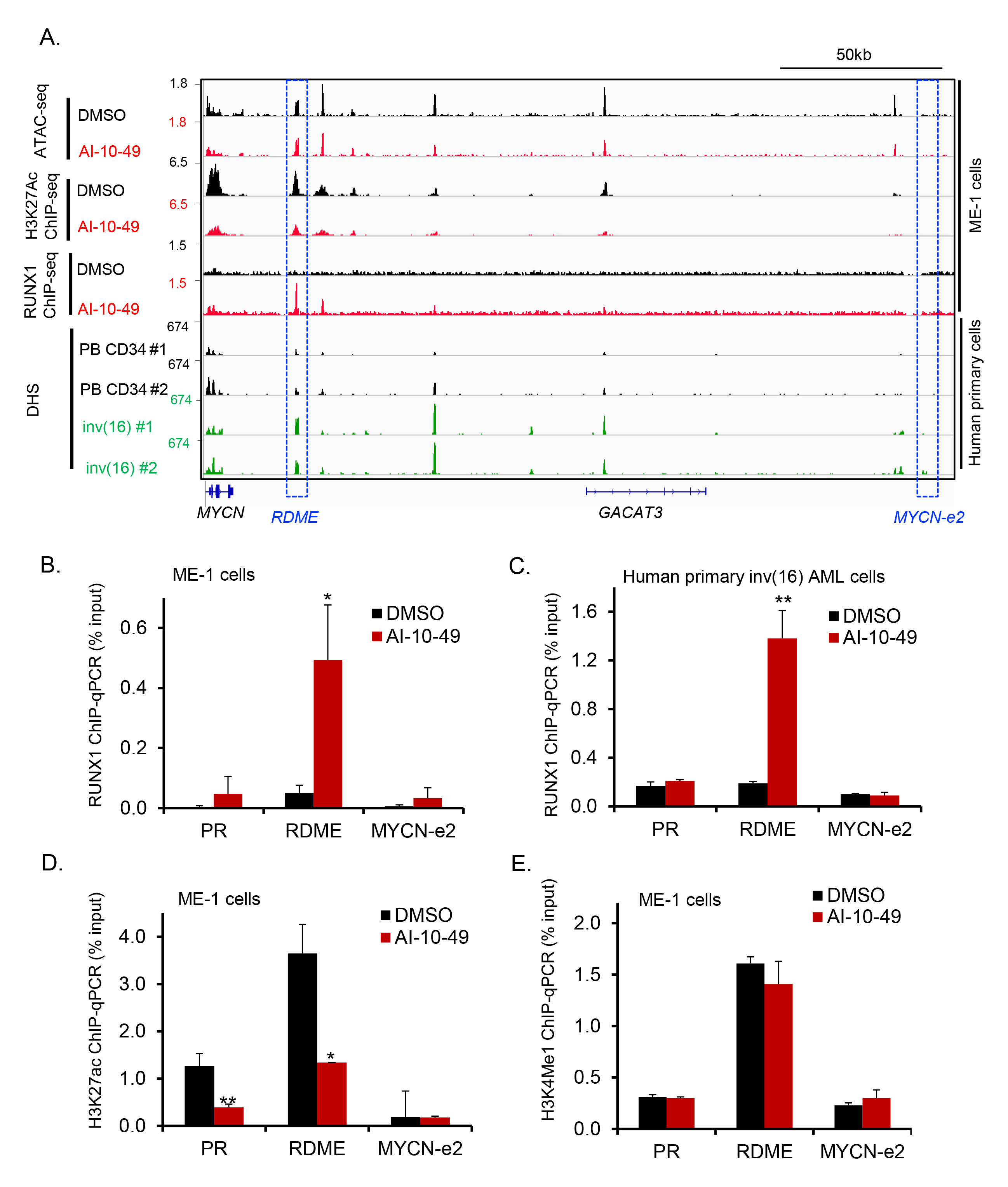
RUNX1 binds to *MYCN* distal enhancer transcriptionally active in inv(16). (**A**). Representative examples of Integrative Genome Viewer (IGV) tracks of ATAC-seq, ChIP-seq, and DNase I-seq analysis in ME-1 cells, healthy peripheral blood CD34+ cells, and purified primary human AML cells. PB CD34: Mobilized CD34+ cells in peripheral blood from healthy individuals; inv(16): CD34+ cells from human primary AML samples with inv(16). (**B-C**). ChIP-qPCR analysis for RUNX1 in DMSO- or AI-10-49-treated cells ME-1 cells (B) and primary human inv(16) AML cells (C). (**D-E**). ChIP-qPCR analysis for H3K27ac (D) and H3K4me1 (E) in DMSO- or AI-10-49 treated ME-1 cells. Histogram representative of triplicate experiments. Error bars represent the SD. Significance was calculated using an unpaired t-test. *p < 0.05 or **p < 0.005.

To further understand the role of *RDME* in inv(16) AML, we analyzed DNase I hypersensitive sites (DHSs) analysis by DNase I-seq, previously reported in primary AML cells. ^22^ We found *RDME* is a highly chromatin-accessible region in primary inv(16) AML cells compared to peripheral blood CD34+ cells from healthy individuals (**Figure 4A**). Furthermore, in addition to inv(16) AML, *RDME* acts as a chromatin-accessible region in multiple AML subtypes such as t(8:21), bi-allelic *CEBPA* mutant, and *RUNX1* mutant AMLs (**Supplemental Figure 4**). Thus, in addition to inv(16) AML, *RDME* may have an essential role in regulating MYCN expression in AML. Taken together, these data suggest *RDME* acts as an oncogenic enhancer in regulating *MYCN* expression in specific subtypes of AML, including inv(16). Furthermore, a recent study identified an enhancer element located 650 kb downstream of *MYCN* TSS is active in specific AML subtypes, ^23^ suggesting distinct *MYCN* enhancers regulate MYCN expression in AML subtype-specific manner. Overall, these results highlight the importance of long-range transcriptional regulation of MYCN in AML.

### *RDME* is required for MYCN transcription and cell survival

To investigate the functional role of *RDME* in inv(16) AML cells, we asked whether deletion of *RDME* by CRISPR/Cas9 approach can affect *MYCN* expression. ME-1 cells were transfected with plasmids expressing Cas9 and two gRNAs for *RDME* and *MYCN-e2* to produce deletions surrounding the RUNX1 binding site in these regions. Our data shows efficient deletion of *RDME* and *MYCN-e2* in ME-1 cells (**Supplemental Figure 5A-F).** *RDME* deletion resulted in a 60% reduction in MYCN transcript levels (**Figure 5A-B**) and a 50% reduction in cell viability due to apoptosis (**Figure 5C**). *RDME* deleted ME-1 cells displayed an increase in the cleaved caspase-3 and cleaved PARP proteins, confirming *RDME* deletion induced apoptosis in ME-1 cells (**Figure 5D**). Meanwhile, the deletion of *MYCN-e2* did not affect MYCN transcript levels or cell viability in ME-1 cells (**Figure 5B-D**). Collectively, our data suggest that *RDME* functions as an oncogenic enhancer to regulate MYCN expression and the viability of inv(16) AML cells.

**Figure 5.**
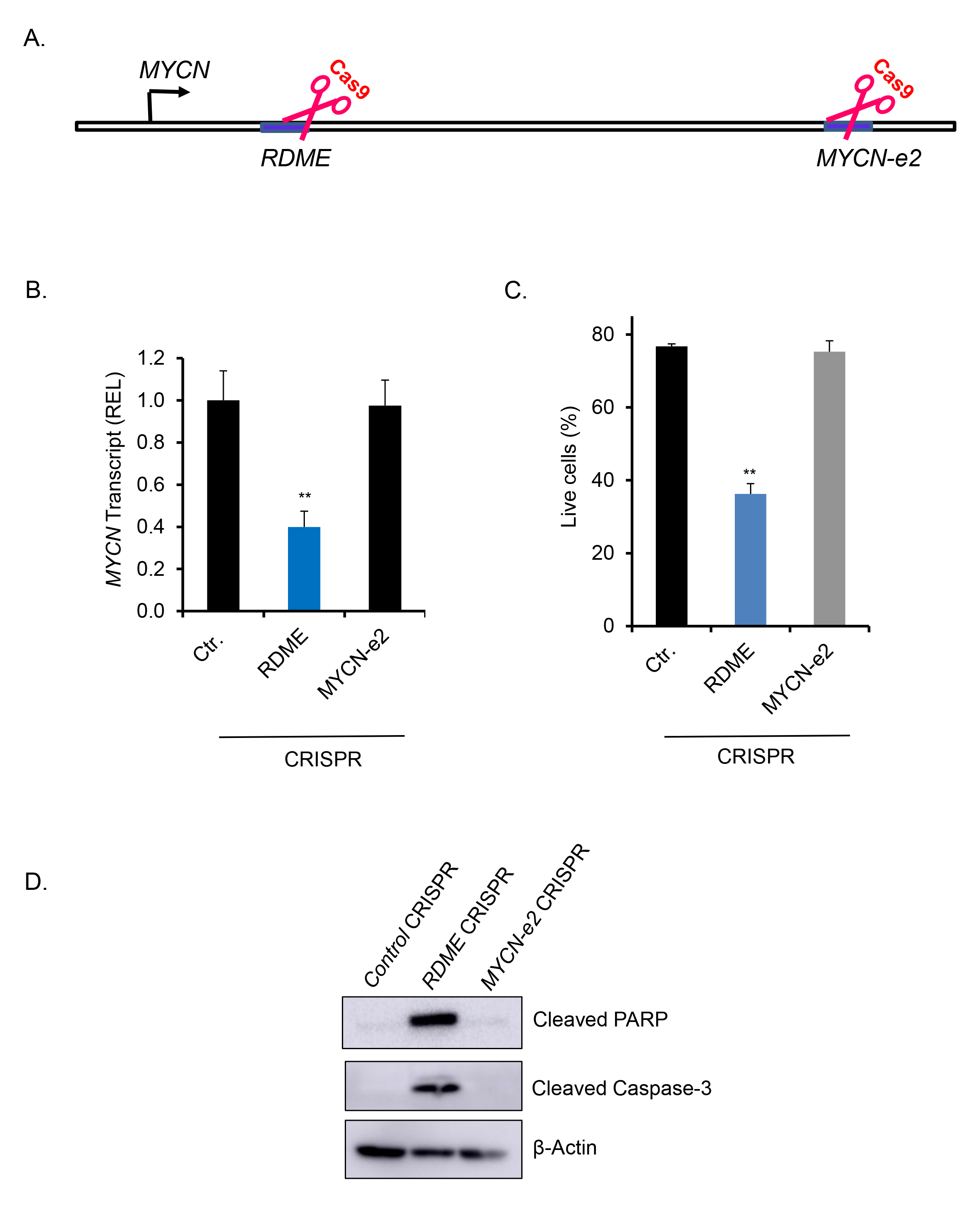
*RDME* is required for N-MYC transcription and cell survival. (**A**). Schematic of CRISPR/Cas9-mediated deletion of *MYCN* enhancer elements. (**B**). N-MYC transcript levels in *MYCN* enhancer deleted ME-1 cells by Real Time RT-PCR. (**C**). Cell survival analysis in control/ *MYCN* enhancer deleted ME-1 cells by Annexin V/7AAD assay. (**D**). Cleaved PARP and cleaved Caspase-3 protein levels in control/ *MYCN* enhancer deleted ME-1 cells by western blot. Histogram representative of triplicate experiments. Error bars represent the SD. Significance was calculated using an unpaired t-test. **p < 0.005.

### *EIF4G1* is a key N-MYC target in inv(16) AML

To identify how N-MYC regulates survival in inv(16) AML cells, we performed CUT&Tag sequencing in ME-1 cells using N-MYC and H3K27ac antibodies (**Figure 6A**). Global comparison of N-MYC CUT&Tag-seq data with H3K27ac CUT&Tag-seq data yielded a strong correlation of N-MYC peak intensity with H3K27ac marks (**Figure 6B**). N-MYC transcriptional targets identified by N-MYC binding and H3K27ac peaks were used to perform gene ontology (GO) analysis. This analysis revealed that N-MYC binding is associated with several pathways, including mRNA metabolism, mRNA splicing, nuclear transport, and chromatin modification (**Figure 6C**). Comparison of N-MYC targets identified by CUT&Tag-seq with previously identified CBFβ-SMMHC transcriptional targets ^14^ and transcripts significantly deregulated in primary human inv(16) AML samples identified Eukaryotic Translation Initiation Factor 4 Gamma 1 (*EIF4G1*) and Kruppel Like Factor 1 (*KLF1*) as potential N-MYC transcriptional targets in inv(16) AML (**Figure 6D**). Of these, only *EIF4G1* transcripts were upregulated in primary human inv(16) AML samples and downregulated with AI-10-49 treatment. *EIF4G1* was one of the regulators of mRNA metabolic process signature genes identified (**Figure 6C**). N-MYC binds to the *EIF4G1* promoter and is associated with a transcriptionally active H3K27ac histone mark (**Figure 6E**). We validated N-MYC binding at *EIF4G1* by conducting ChIP-qPCR (**Figure 6F**). DNase I hypersensitive site analysis ^22^ revealed that the *EIF4G1* promoter is a highly accessible chromatin region in primary inv(16) AML cells compared to peripheral blood CD34+ cells from healthy individuals (**Figure 6E**). *EIF4G1* transcript levels were upregulated in primary inv(16) AML patient samples (**Figure 6G-H**). We observed downregulation of eIF4G1 at mRNA and protein levels with AI-10-49 in inv(16) AML cells (**Supplemental Figure 6**). Genomic deletion of *MYCN* in ME-1 cells by CRISPR/Cas9 RNPs resulted in a marked reduction of eIF4G1 protein levels (**Figure 6I**). Taken together, *EIF4G1* is a key N-MYC target with potential roles in leukemic cell survival in inv(16) AML.

**Figure 6.**
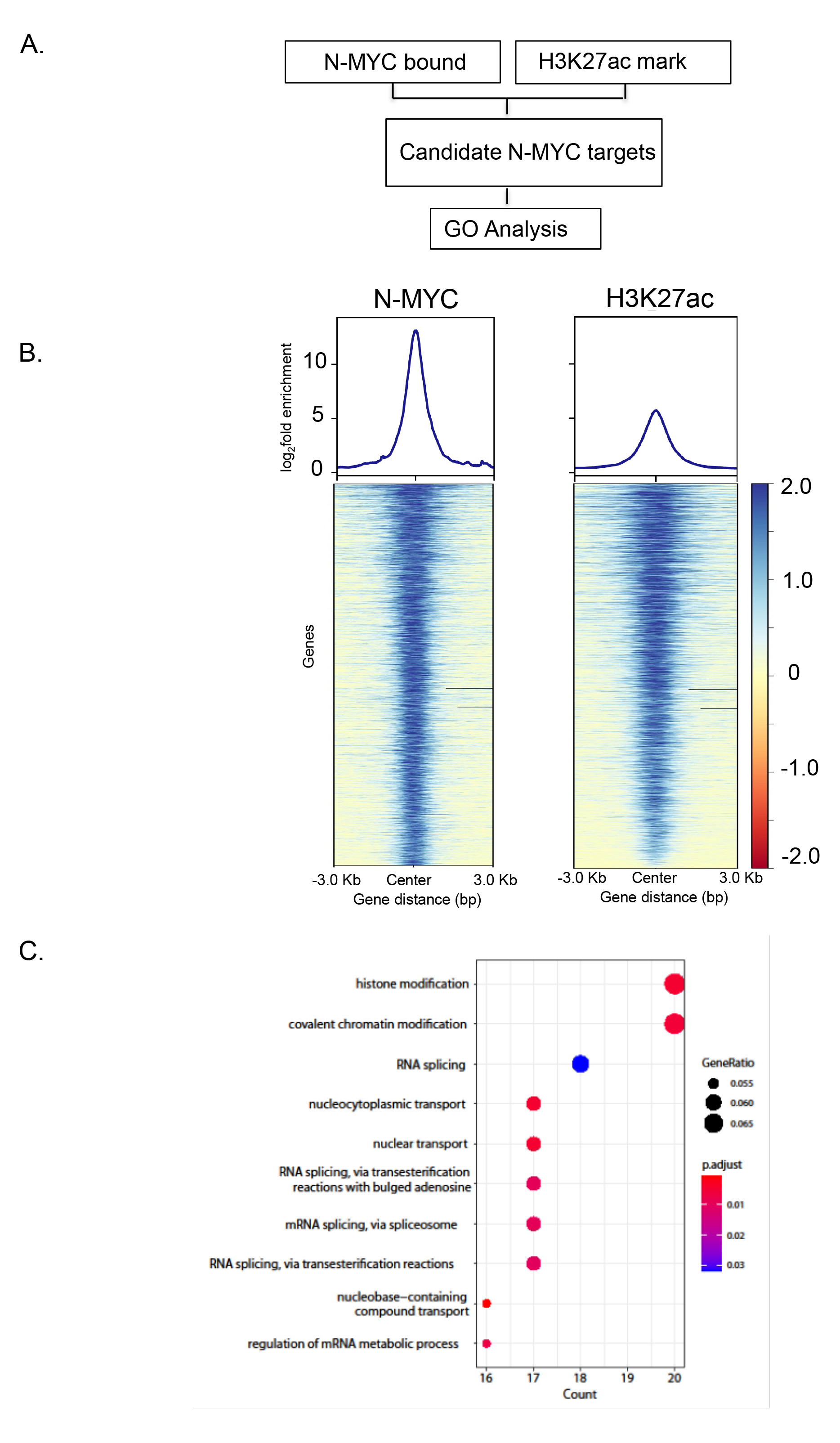
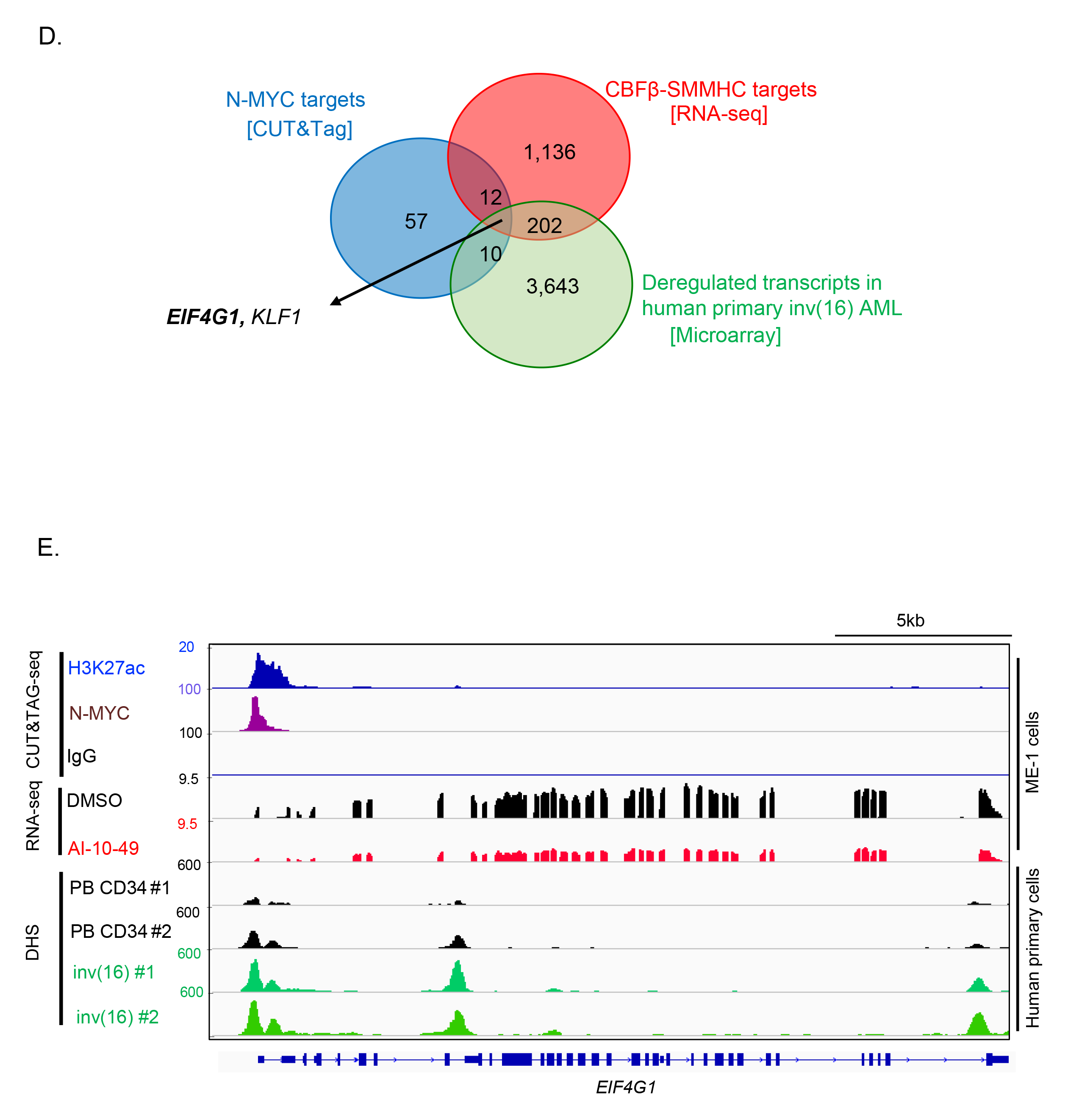
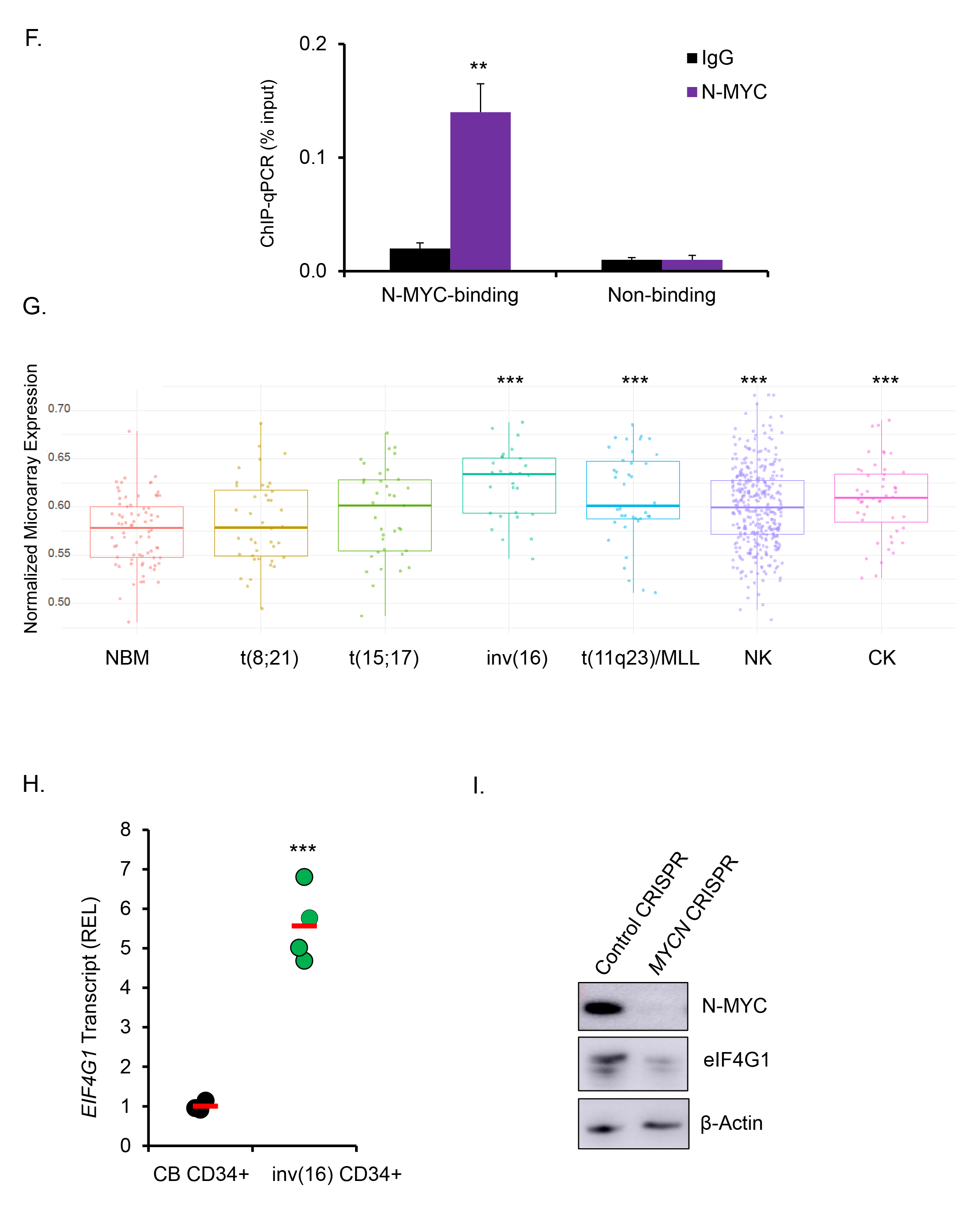
eIF4G1 is a key N-MYC target in inv(16) AML. (**A**). Filtering scheme for identifying N-MYC targets in inv(16) AML. N-MYC -bound regions (p < 0.05) were intersected with the H3K27ac histone mark (p < 0.05). Candidate MYCN targets identified were used for gene ontology enrichment analysis. (**B**). Genome-wide distribution of N-MYC and H3K27ac peaks in CUT&TAG-seq. (**C**). Dot plot of gene ontology enrichment analysis of N-MYC targets identified in inv(16) AML. The diameter indicates the number of genes overlapping the gene ontology term, and the color indicates the enrichment P-value. (**D**). Venn diagram showing the overlap between significant differentially expressed genes in AI-10-49 treated ME-1 cells (±2 fold change; adj. *p* < 0.05) identified in the RNA-seq, transcripts significantly deregulated in human inv(16) AML CD34+ cells compared to normal bone marrow CD34+ cells in Haferlach dataset extracted from Leukemia Gene Atlas (expression array) database and peaks called in N-MYC CUT&Tag. (**E**). Representative examples of Integrative Genome Viewer (IGV) tracks of CUT&Tag-seq, RNA-seq, and DNase I-seq analysis in ME-1 cells, healthy peripheral blood CD34+ cells, and purified human primary AML cells. (**F**). ChIP-qPCR analysis for N-MYC binding region in *EIF4G1* promoter and a non-binding region in *EIF4G*1 locus in ME-1 cells. Histogram representative of triplicate experiments. (**G**). Normalized expression of EIF4G1 in Haferlach dataset extracted from Leukemia Gene Atlas (expression array) databases. (**H**). eIF4G1 transcript levels in human cord blood CD34+ cells and human primary inv(16) AML CD34+ cells. Each symbol represents the average of a triplicate experiment from one sample, and the average value of the group is shown in red. (**I**) ME-1 cells were transfected with Cas9 and control gRNAs/ *MYCN* gRNAs [Synthego Gene Knockout kit V2] by RNP approach and analyzed N-MYC and eIF4G1 protein levels by western blot. Error bars represent the SD. Significance was calculated using an unpaired t-test. **p < 0.005 or ***p < 0.0005.

c-MYC and N-MYC control their expression by repressing each other at promoter sites, thereby acting in autoregulatory feedback loops in neuroblastoma cells ^24,25^ Deletion of N-Myc in steady state hematopoiesis does not affect c-Myc expression. ^26^ To address the interplay between N-MYC and c-MYC in inv(16) AML cells, we asked whether N-MYC regulates c-MYC expression. Our data suggest that N-MYC binds to *MYC* promoter transcriptionally active in primary human inv(16) AML cells. (**Supplemental Figure 7A, B).** Genomic deletion of *MYCN* in ME-1 cells by the CRISPR/Cas9 approach resulted in a marked reduction of *MYC* transcript levels (**Supplemental Figure 7C).** Taken together, N-MYC positively regulates MYC expression in inv(16) AML cells.

### eIF4G1 is required for inv(16) AML maintenance

Since N-MYC plays an essential role in inv(16) leukemia maintenance and *EIF4G1* is a major target of N-MYC, we reasoned that eIF4G1 would be instrumental in regulating inv(16) AML survival. To test this, we applied CRISPR/Cas9 technology using RNPs to delete *EIF4G1* in inv(16) AML cells. ME-1 cells transfected with Cas9 protein and three *EIF4G1* CM-gRNAs produced efficient deletion of *EIF4G1* genomic regions (**Supplemental Figure 7A-C)**, resulting in a 95% reduction in eIF4G1 protein levels (**Figure 7A**). *EIF4G1* deletion in ME-1 cells significantly lowered cell viability due to apoptosis (**Figure 7B**). The onset of apoptosis was accompanied by the cleavage of caspase 3 and cleaved PARP proteins in ME-1 cells (**Figure 7C**). To further assess the role of eIF4G1 in inv(16) AML cell survival, we conducted *EIF4G1* genomic editing in primary human inv(16) AML cells, as described in **Figure 3A**. T cell-depleted primary inv(16) AML cells were transfected with Cas9 protein and three *EIF4G1* gRNAs resulting in an 85% reduction in eIF4G1 protein levels (**Figure 7D**). *EIF4G1* deletion significantly reduced the clonogenic potential of primary inv(16) AML cells, as reflected by a marked decrease in colony number (**Figure 7E**). The requirement of eIF4G1 in inv(16) leukemia in vivo was tested by transplanting the edited leukemic cells into irradiated (280 cGy) NSGS mice via tail vein injection. Engraftment efficiency of control and *EIF4G1* edited leukemic cells in the bone marrow five days after transplantation was similar between groups (**Supplemental Figure 7D**). AML burden estimated by the frequency of hCD45+ hCD33+ AML stem/progenitor cells was significantly reduced in the bone marrow and peripheral blood of mice transplanted with *EIF4G1* edited AML cells compared to control mice (**Figure 7F-H**), suggesting that eIF4G1 is required for the maintenance of inv(16) leukemic cells. In addition, the median leukemic latency in mice transplanted with *EIF4G1* deleted AML cells was significantly extended from 72 days (control group) to 108 days (*EIF4G1* deleted group; p <0.05) (**Figure 7I**). Collectively, these results demonstrate that *EIF4G1* is a critical N-MYC target instrumental in inv(16) AML maintenance.

**Figure 7.**
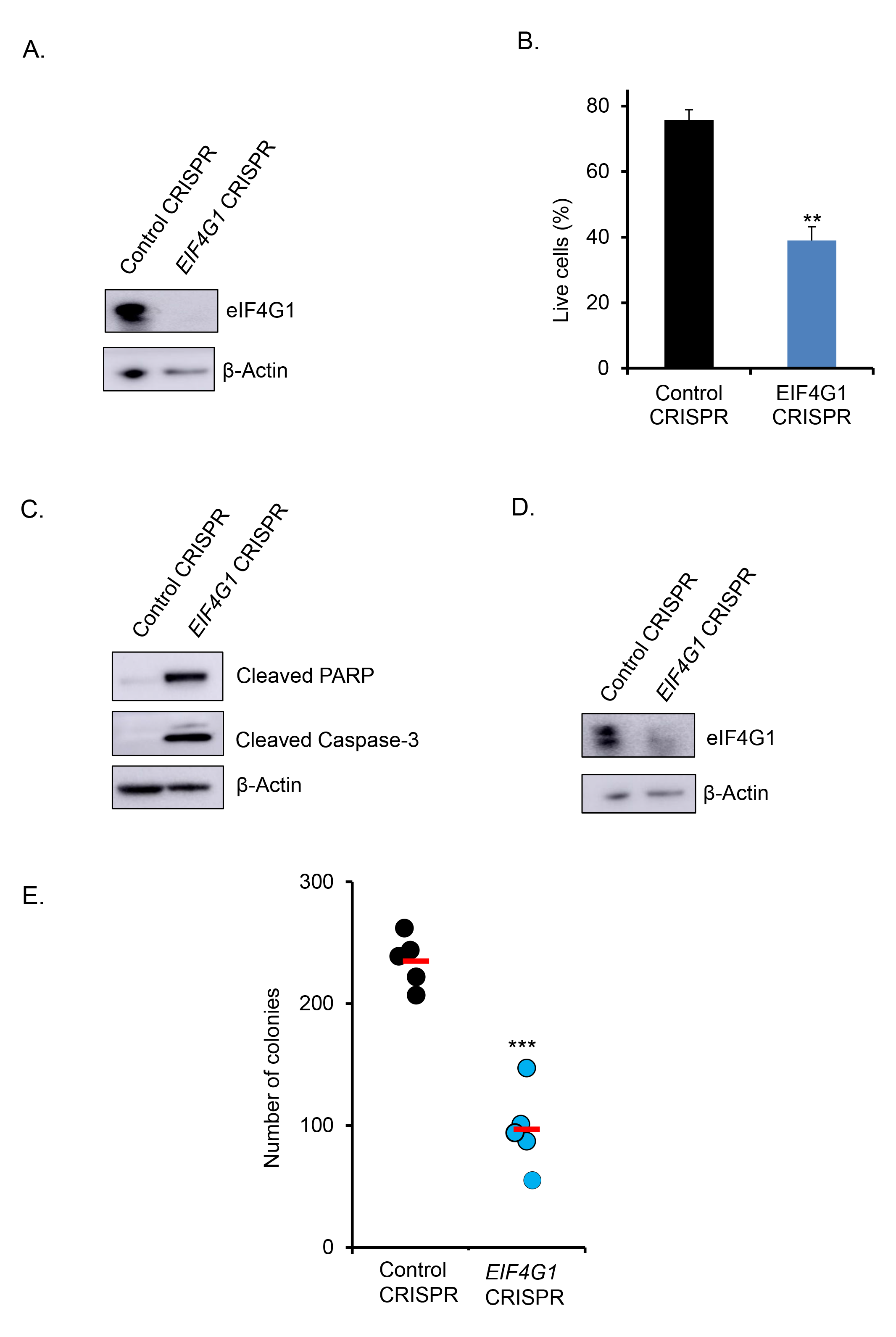
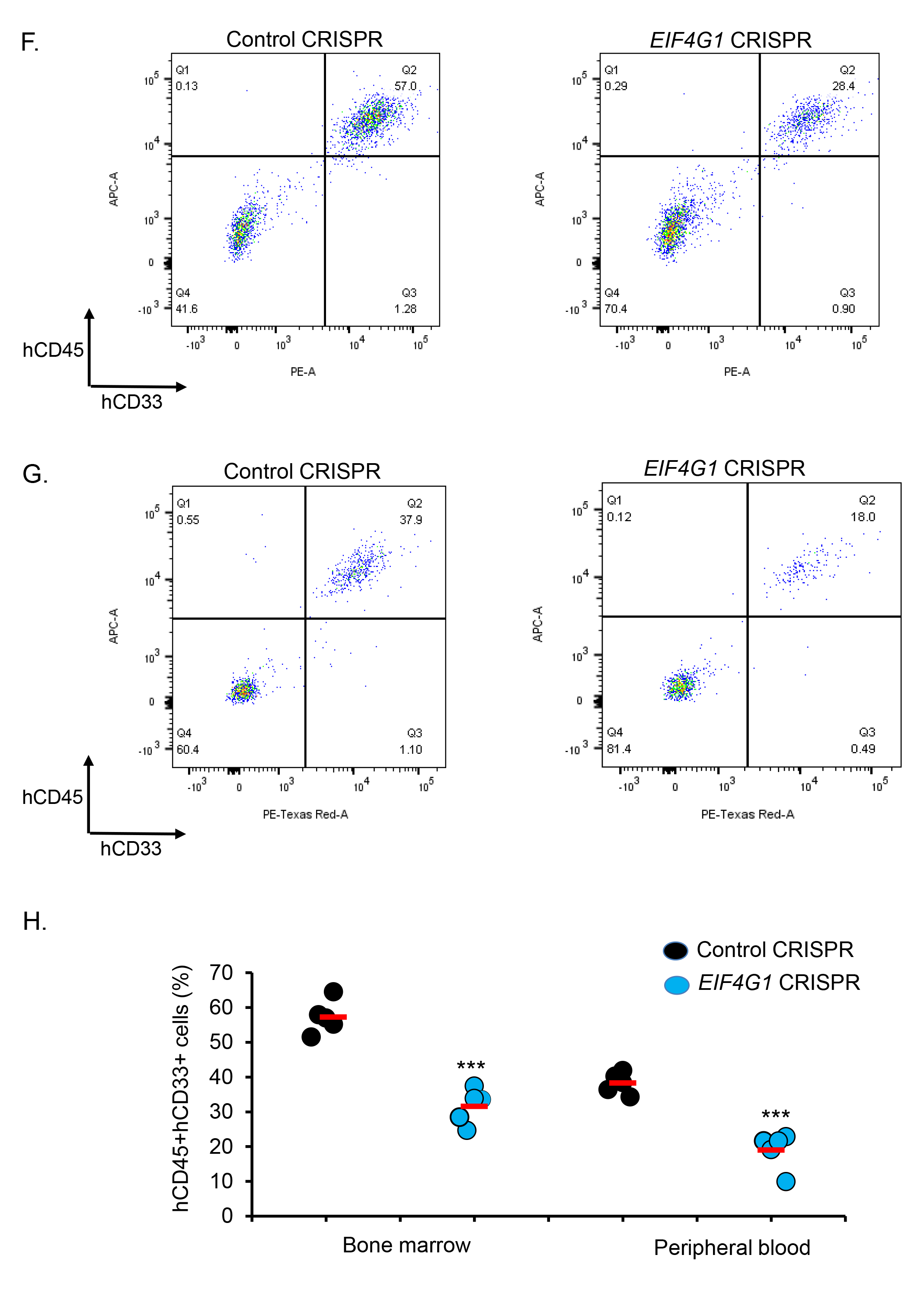
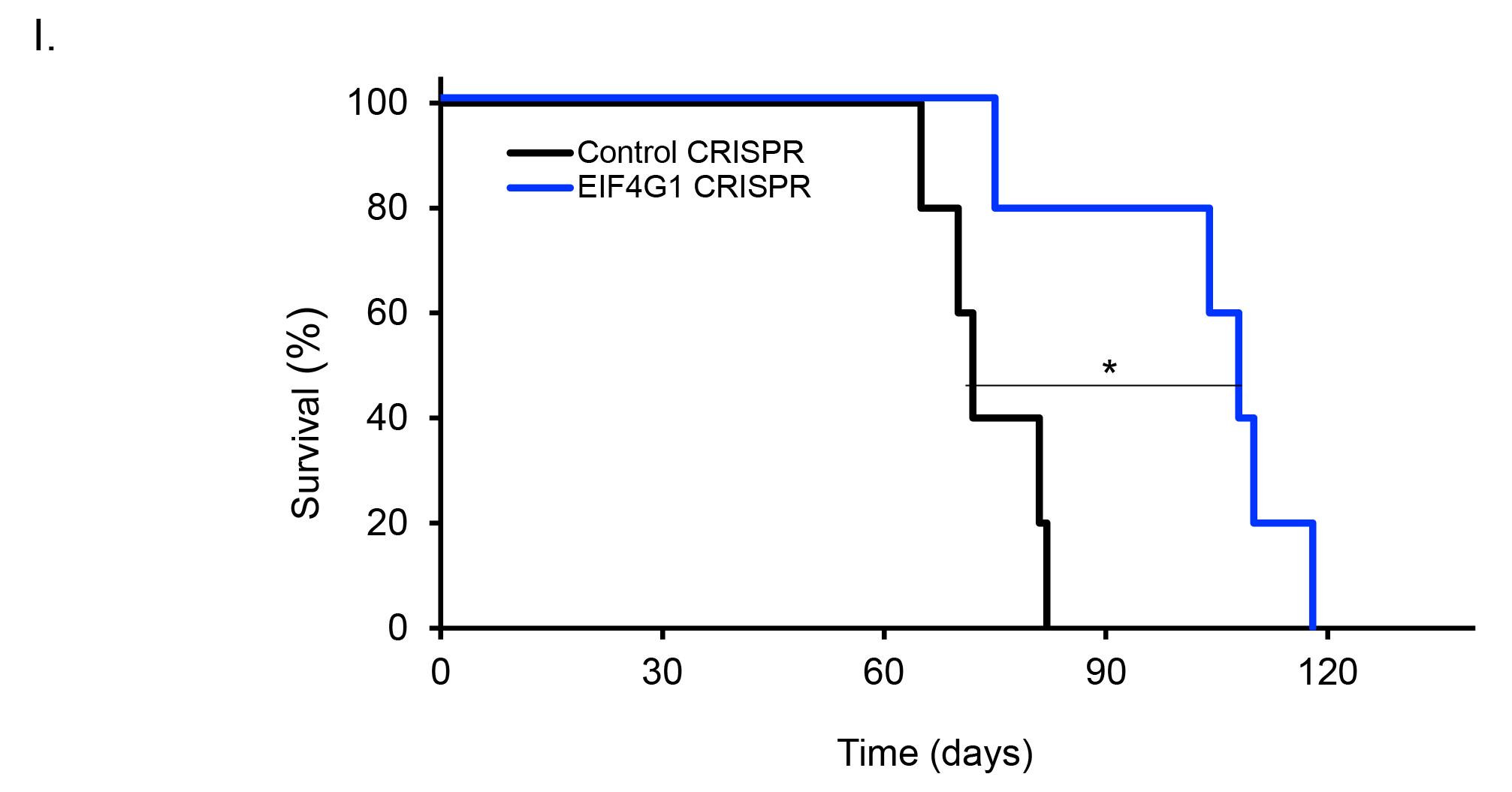
eIF4G1 is required for inv(16) AML maintenance. (**A**). ME-1 cells were transfected with Cas9 and control gRNA/ pool of 3 *EIF4G1* gRNAs by RNP approach and analyzed eIF4G1 protein levels by western blot. (**B**). Cell survival analysis in control/ *EIF4G1* edited ME-1 cells by Annexin V/7AAD assay. Histogram representative of triplicate experiments. (**C**). Cleaved PARP and cleaved Caspase-3 protein levels in control/ *EIF4G1* edited ME-1 cells by western blot. (**D**). Human primary inv(16) AML cells were transfected with Cas9 and control gRNA/ pool of three *EIF4G1* gRNAs by RNP approach and analyzed eIF4G1 protein levels by western blot. (**E**). Colony counts for methylcellulose colony-forming assay performed upon 12 days of control/ *EIF4G1* edited human primary inv(16) AML cells. Data representative of the four replicates. The average value of each group is shown in red. (**F-G**). Representative flow cytometry plots showing gating and frequency of hCD45+ hCD33+ cells in bone marrow (F) and peripheral blood (G) of NSGS mice transplanted with control/ *EIF4G1* edited primary inv(16) AML cells five weeks after transplantation. (**H**). Flow cytometric quantification of hCD45+ hCD33+ cells in NSGS mice transplanted with control/ *EIF4G1* edited primary inv(16) AML cells five weeks after transplantation. Each symbol represents a mouse. The average value of each group is shown in red. (**I**). Kaplan-Meier survival curve of NSGS mice transplanted with control/ *EIF4G1* edited primary inv(16) AML cells (n=5/group). Error bars represent the SD. Significance was calculated using an unpaired t-test (B, E, H) and log-rank test (I). *p < 0.05 or **p < 0.005 or ***p < 0.0005.

## Discussion

Both c-Myc and N-Myc are essential for normal development, as shown by embryonic lethality at mid-gestation due to the lack of either gene in mouse models. ^27^ N-Myc is vital for fetal HSC proliferation. ^28^ Deletion of *c-Myc* in adult mouse hematopoiesis results in the accumulation of HSCs that are defective in further differentiation. ^29,30^ Meanwhile, deletion of *N-Myc* in adult mouse hematopoiesis did not display any phenotype. ^26^ Combined deletion of both *c-Myc* and *N-Myc* in adult mice results in pancytopenia and rapid lethality, demonstrating that both c-Myc and N-Myc are required for HSC proliferation, self-renewal, and survival. ^26^ Thus, c-Myc and N-Myc proteins have redundant functions in normal hematopoiesis. The oncogenic role of c-MYC is well established in AML. ^14,31,32^ While the requirement of c-MYC and the pathways orchestrated by c-MYC are widely investigated in hematological malignancies, the role of N-MYC in leukemia remains largely unexplored.

In this study, utilizing primary human inv(16) AML samples and a patient-derived xenograft model for inv(16) AML, we revealed a previously unidentified role of N-MYC in leukemogenesis. The oncogenic function of N-MYC has been widely studied in neuroblastoma, and emerging studies show its involvement in prostate cancer, pancreatic cancer, and retinoblastoma. ^33^ *MYCN* amplification is reported in AML and lymphoma. ^34,35^ N-MYC overexpression is reported in pediatric AML, erythroleukemia, and Burkitt lymphoma. ^36-38^ Overexpression of *Mycn* in mouse bone marrow cells results in enhanced proliferation and self-renewal of myeloid progenitors resulting in AML development. ^39^ T-cell progenitor-specific N-Myc overexpression in mice induces peripheral T-cell lymphoma (PTCL). ^40^ Here, we demonstrate that CBFβ–SMMHC oncogene in inv(16) AML cells modulates N-MYC transcript levels, and N-MYC is a critical regulator of inv(16) AML cell survival.

Both c-Myc and N-Myc are expressed in HSCs. ^26,41^ No compensatory upregulation of *c-Myc* is observed upon loss of *N-Myc*, suggesting that when *N-Myc* is deleted, the unchanged *c-Myc* transcript levels can provide HSC survival and proliferation. ^26^ Having shown that c-MYC is required for inv(16) AML cell survival, ^14^ how can we explain the loss of N-MYC regulating survival of inv(16) AML cells? Multiple explanations can be made to describe the oncogenic dependency of N-MYC in inv(16) AML cells. c-MYC and N-MYC can regulate their expression via autoregulatory feedback loops in neuroblastoma cells. ^24,25,42^ We observed that N-MYC binds to the *c-MYC* promoter in inv(16) AML cells and is associated with MYC transcription (**Supplemental Figure 7**), suggesting N-MYC may have essential roles in c-MYC regulated inv(16) AML cell survival reported before. ^14^ Thus, even though N-MYC does not regulate c-MYC expression in normal HSCs, N-MYC positively regulates c-MYC expression in AML cells, identifying fundamental differences in the interplay between c-MYC and N-MYC in normal and leukemic hematopoiesis. Secondly, even though the N-Myc protein functions very similarly to c-Myc, ^43^ there are significant differences in the expression pattern of c-Myc and N-Myc proteins in mouse hematopoietic compartments. The expression of c-Myc and N-Myc proteins in HSC is mutually exclusive. ^41^ While N-Myc is expressed at high levels in self-renewing and quiescent HSCs, c-Myc is expressed in multiple progenitor compartments. In addition, N-Myc overexpression can reprogram mature blood cells to HSCs. Thus, the unique expression pattern of c-Myc and N-Myc could have distinct roles in AML cell survival. Thirdly, c-Myc and N-Myc proteins may have different binding partners, thereby exerting distinct transcriptional programs in AML cells. c-Myc and N-Myc proteins differ in MIZ1 binding and have distinct functions in medulloblastoma. ^44^ c-Myc and N-Myc protein-protein interactions in AML cells remain largely unknown. Thus, N-MYC and c-MYC may have overlapping and distinct transcriptional targets in AML. Further studies addressing how N-MYC functions differently from c-MYC in AML initiation and maintenance warrant novel insights.

We identified *EIF4G1* as a key N-MYC target regulating inv(16) AML cell survival. eIF4G1 plays a vital role in translation initiation by acting as an adapter that enhances the assembly of the eIF4F complex by recruiting eIF4E and eIF4A. Cancer cells are highly dependent on increased eIF4F activity compared to normal cells. ^45,46^ eIF4E is overexpressed and known to have oncogenic potential in M4 and M5 subtypes of AML. ^47-49^ eIF4A is overexpressed in AML, and its inhibition can induce anti-leukemic effects in FLT3-ITD AML. ^50^ eIF4G1 expression is highly activated in breast, prostate, and squamous cell lung cancers and is associated with metastatic progression and inferior survival. ^51-53^ Neuroblastoma patients with *MYCN* amplification present elevated *eIF4G1* transcript levels and are associated with significantly lower survival. ^54^ To our knowledge, this is the first report to show the oncogenic role of eIF4G1 in AML. While previous studies have implicated that eIF4G1 acts as an oncogene by exerting its role in translation initiation, we found that eIF4G1 plays a role in mRNA metabolism in inv(16) AML cells. eIF4G1 plays a major role in mRNA metabolism by degradation of specific nuclear mRNAs. ^55^ New evidence suggests that altered mRNA metabolism plays a pivotal role in leukemia. ^56,57^ Although gene expression profiling in inv(16) AML has been associated with alterations in mRNA metabolism, ^58^ a mechanistic explanation for deregulation in mRNA metabolism in inv(16) leukemogenesis remains unknown. Further experiments will be required to identify the specific role of eIF4G1 in mRNA metabolism and AML cell survival.

Our results on the oncogenic role of the N-MYC/ eIF4G1 axis provide key insights for developing better therapeutic strategies for patients with inv(16) AML. The development of bromodomain inhibitors has gained wide attention in indirectly inhibiting c-MYC and N-MYC in cancer. ^16,17^ However, recent clinical trials revealed that most of these inhibitors had limited clinical utility due to unexpected toxicities. ^59^ So, a promising alternative is to target c-MYC and N-MYC downstream pathways in AML. Since c-MYC and N-MYC play a major role in normal hematopoiesis, targeting pathways activated by c-MYC and N-MYC in AML cells will have major therapeutic benefits. The recent development of SBI-756, an eIF4G1 small molecule inhibitor, has demonstrated promising efficacy in multiple cancer types, including melanoma and B-acute lymphocytic leukemia. ^60,61^ Future studies using combinations of AI-10-49 with SBI-756 should lead to greater therapeutic responses for inv(16) AML.

In conclusion, we have discovered N-MYC as a major leukemic driver in inv(16) AML. Furthermore, we also identified eIF4G1 as a key N-MYC transcriptional target in AML survival, providing novel insights on developing improved treatment avenues for patients with inv(16) AML.

## Methods

### Mice

All animal experiments were performed in accordance with a protocol reviewed and approved by the Medical College of Wisconsin Institutional Animal Care and Use Committee. NSGS [NOD/SCID-IL2RG–SGM3] mice have been previously described^18,19^ and maintained at the Medical College of Wisconsin animal facility.

### Cell Line and Primary Hematopoietic Cell Cultures

Human inv(16) AML ME-1 cells were cultured in RPMI 1640 with 20% fetal bovine serum, 25 mM HEPES, 100 U/mL Penicillin, 100 μg/mL Streptomycin and 1 μl/mL Plasmocin. Human cord blood samples were collected from the Medical College of Wisconsin Tissue bank, and CD34+ cells were isolated using the CD34 Microbead kit (Miltenyi Biotec, #130-046-702). The use of the cord blood samples for research purposes was approved by the Ethics Committee of the Medical College of Wisconsin. Human AML samples were received from AML banks by Martin Carroll (University of Pennsylvania) and Carsten Mueller Tidow (University Hospital, Heidelberg, Germany). Hematopoietic CD34+ cells were isolated from human AML bone marrow samples using the CD34 Microbead kit. All patients gave written consent for the use of their samples. Personal information from AML and cord blood samples was unavailable as the samples were anonymized. Human cord blood CD34+ cells, as well as human primary leukemic cells, were cultured in StemSpan SFEM II (STEMCELL Technologies, #09605), 100 U/mL Penicillin and 100 μg/mL Streptomycin supplemented with 10 ng/mL human recombinant TPO, 10 ng/mL human recombinant FLT3L (10 ng/mL), 100 ng/mL human recombinant SCF, 10 ng/mL human recombinant IL3, and 20 ng/mL human recombinant IL-6. Cell cultures were routinely tested for mycoplasma contamination using Universal Mycoplasma Detection Kit (ATCC, #30-1012K) and PlasmoTest Mycoplasma Detection Kit (Invivogen, #rep-pt1).

### Small molecule inhibitors and cytokines

AI-10-49 and JQ1 were purchased from ApexBio. All cytokines were purchased from Peprotech.

### CRISPR/Cas9 genomic editing of *MYCN* enhancers

The sgRNAs specific for 5’ to the region of interest were cloned in pLentiCRISPRv2 (Addgene, #52961). sgRNAs corresponding to 3’ to the region of interest was cloned in pDecko-mCherry (Addgene, #78534). The puromycin resistance cassette in pLentiCRISPRv2 was replaced by a GFP gene using standard cloning techniques. Oligonucleotide sequences are listed in Table S1. 2×10E6 ME-1 cells were nucleofected with CRISPR/Cas9 plasmids (2 μg each) using Nucleofector Technology (Lonza Biologics) with the program X-01 and Amaxa Cell Line Nucleofector Kit V. Samples were sorted by flow cytometry 24 hr later. Cells were cultured overnight, and dead cells were eliminated by dead cell removal kit (Miltenyi Biotec, #130-090-101).

### CRISPR/Cas9 genomic editing of *MYCN* and *EIF4G1*

Gene editing experiments in primary human inv(16) AML CD34+ cells were conducted using the ribonucleoprotein (RNP) complex. Gene Knockout Kit v2 (Synthego) consisting of three chemically modified Guide RNAs (CM-sgRNAs) was used to target *MYCN* and *EIF4G1* in ME-1 cells and primary human AML CD34+ cells. Negative Control Scrambled sgRNA and SpCas9 protein were purchased from Synthego. To assemble the RNP complex, sgRNAs were combined with Cas9 protein and incubated for 10 minutes at room temperature (RT).

Primary human inv(16) AML CD34+ cells were T-cell depleted using CD3 Microbeads (Miltenyi Biotec, #130-050-101) and nucleofected with the RNP complex, composed of 100 pmols of Cas9 and 300 pmols of CM-sgRNA, using the P3 Primary Cell 4D-Nucleofector X Kit for Amaxa 4-D device (Lonza, #V4XP-3032). 2×10E5 cells per condition were nucleofected in separated strip wells using program EO-100. Cell viability was assessed by flow cytometry 24 hours post-nucleofection and gene editing efficacy was evaluated 72 hours post-nucleofection by Inference of CRISPR editing (ICE) and western blot analysis.

### CRISPR/Cas9-induced gene editing analysis by Inference of CRISPR Editing (ICE)

Genomic DNA was isolated from edited cells 24 hours after nucleofection. The edited region was PCR amplified, and the PCR product was sequenced by the Sanger method. The chromatograms were analyzed by ICE (https://ice.synthego.com). The percentage of editing was calculated according to the frameshift produced in the edited chromatogram compared to the control sequence. Primers used in these PCRs are listed in Table S1.

### Colony Forming Unit Assay

Twenty-four hours after nucleofection of AML inv(16) CD34+ cells with the RNP complex, cells were resuspended in MethoCult Express (STEMCELL Technologies, #04437), plated at 3,000-5,000 cells/mL on a 12-well plate and cultured for 12 days to form colonies.

### Transplantation of Gene-edited AML cells in NSGS mice

Twenty-four hours after nucleofection of AML inv(16) CD34+ cells with the RNP complex, 1×10E6 cells were transplanted into sub-lethally (280 rads) irradiated six to eight-week-old immunodeficient NSGS mice (n = 5 per group) by tail vein injection. Successful engraftment of edited AML cells was evaluated by flow cytometry of bone marrow aspirates five days later. The leukemic burden was analyzed by flow cytometry of peripheral blood and bone marrow aspirates five weeks later. Mice were sacrificed after visible characteristics of AML, including reduced motility and grooming activity, hunched back, and pale paws (anemia).

### Flow Cytometry

For flow cytometry, cells were washed twice with 1% BSA in PBS, stained for 20-30 min at 4C in the dark, and analyzed with a BD LSRII flow cytometer. Flow cytometry analysis was performed using FlowJo Software. The following antibodies were used: Human CD33 Monoclonal Antibody (eBioscience, # 12-0338-42), Human CD45 Monoclonal Antibody (BD Biosciences, #555485), CD11b Monoclonal Antibody (BD Biosciences, #555388), and CD15 Monoclonal Antibody (BD Biosciences, # 555401).

### Annexin V Assay

For testing the role of *MYCN* and *eIF4G1* silencing in inv(16) cell survival, ME-1 cells were transfected with control / *MYCN/ EIF4G1* CM-sgRNAs and assessed live cells (7AAD-Annexin V-) by Annexin V assay. Annexin assay was conducted 14 days after nucleofection. For testing the role of *MYCN* enhancer deletion in inv(16) AML cell survival, ME-1 cells were transfected with empty vector/Cas9 (Ctr.) or *MYCN* enhancer sgRNA/Cas9 constructs, sorted cells 24 hr later and assessed live cells (7AAD-Annexin V-) by Annexin V assay 14 days later. For the detection of apoptotic cell death, the Annexin V Apoptosis Detection Kit I (BD Bioscience, #559763) was used as per the manufacturer’s instructions. Briefly, cells were centrifuged at 2000 rpm for 10 min, resuspended in 100 μl 1X Annexin V binding buffer, added 5 μl Annexin-PE and 10 μl 7AAD and incubated for 15 min at RT in the dark, followed by adding 500 μl 1X Annexin binding buffer. Cell viability was determined as the percent of 7-AAD negative/ Annexin V negative cells with a BD LSRII flow cytometer.

### Quantitative RT-PCR Analysis

To evaluate MYC, MYCN, and EIF4G1 transcriptional regulation by AI-10-49 and JQ1 in inv(16) cells, ME-1 cells were treated with AI-10-49/ JQ1 for corresponding time points. mRNA was isolated from three independent experiments and conducted qRT-PCR. For testing the role of *MYCN* enhancer deletion in MYCN transcriptional regulation in inv(16) cells, ME-1 cells were transfected with empty vector/Cas9 (Ctr.) or MYCN enhancer sgRNA/Cas9, sorted cells 48 hrs later, and isolated mRNA from three independent experiments 24 hr later. MYCN expression was estimated by qRT-PCR analysis. Total mRNA was isolated with a PureLink RNA Mini Kit (life technologies) and cDNA synthesis was performed with a SuperScript III kit (Life Technologies), as per the manufacturers’ instructions. Quantitative PCR analysis was conducted on an Applied Biosystems QuantStudio™ 6 Flex Real-Time PCR System with Power SYBR Green PCR Master Mix (Applied Biosystems). Expression levels were determined with the delta Ct method and normalized to GAPDH mRNA. Sequences of primers are provided in Table S1

### Chromatin Immunoprecipitation (ChIP) qPCR

ME-1 cells were treated with DMSO or AI-10-49 (1 μM) for 6 hrs. Cross-linking of proteins to DNA was accomplished by adding 1% formaldehyde for 10 min to cultured cells at RT. After neutralization with glycine, cells were lysed in a lysis buffer with protease inhibitors, and samples were sonicated to an average DNA length of 200–400 bp with a bioruptor (Diagenode). After sonication, the chromatin was immunoprecipitated with 5-10 μg of the antibody of interest at 4C overnight. Antibody bound complexes were isolated with Dynabeads (Life Technologies). DNA was purified using the phenol-chloroform isoamyl-alcohol method. Immunoprecipitated DNA was analyzed by qPCR on a QuantStudio™ 6 Flex Real-Time PCR System (Applied Biosystems) with Power SYBR Green PCR Master Mix and calculated as % of input. For ChIP qPCR in human primary AML sample with inv(16), CD34+ cells were enriched using CD34 MicroBead Kit (Miltenyi Biotec, #130-046-702) and cultured overnight, followed by dead cell removal by dead cell removal kit (Miltenyi Biotec, #130-090-101). Cells were treated with DMSO/ AI-10-49 (5 µM) for 8 hrs, followed by the ChIP procedure mentioned above. Details of the sequences of primers are provided in Table S1. The following antibodies were used: RUNX1 polyclonal (Abcam, #ab23980), H3K4Me1 polyclonal (Abcam, #ab8895), H3K27ac polyclonal (Abcam, #ab4729), N-Myc (D1V2A) Monoclonal Antibody (Cell Signaling Technology, #84406) and normal Rabbit IgG (Cell Signaling Technology, #2729).

### Cleavage Under Targets and Tagmentation (CUT&Tag)

CUT&Tag was performed according to the protocol reported by the Henikoff laboratory.^62^ ME-1 cells were harvested and centrifuged at 600×*g* for 3 min at RT. 0.5×10E6 aliquots of cells were washed once in 1.5ml of Wash Buffer (20 mM HEPES pH 7.5; 150 mM NaCl; 0.5 mM Spermidine; 1× Protease inhibitor cocktail) by gentle pipetting and resuspended in 100 µl Wash buffer. Bio-Mag Plus Concanavalin A coated beads (Polysciences) were prepared by washing twice in 1.5ml of binding buffer (20 mM HEPES pH 7.9, 10 mM KCl, 1mM CaCl_2_ and 1mM MnCl_2_) and resuspended in binding buffer (10 µl per sample). 10 µl of activated beads were added per sample with gentle vortexing and placed on an end-over-end rotator for 10 min at RT. The unbound supernatant was removed by placing the tube on the magnetic stand, and bead bound cells were resuspended in 50µl of ice-cold Dig-wash buffer (20 mM HEPES pH 7.5; 150 mM NaCl; 0.5 mM Spermidine; 1× Protease inhibitor cocktail; 0.05% Digitonin) containing 2 mM EDTA, 0.1% BSA and a 1:50 dilution of the appropriate Primary antibody (N-Myc, H3K27Ac or IgG). After overnight incubation at 4 °C, the primary antibody was removed, and the cells were resuspended in 100 µl Dig-wash Buffer containing 1:100 dilution secondary antibody. Cells were incubated at RT for 45 min and washed twice in 1 ml Dig-Wash buffer to remove unbound antibodies. The pA-Tn5 adapter complex containing 2.5 µl of 20x CUTANA™ pAG-Tn5 pre-loaded adapter complex (EpiCypher, #15-1017) was prepared in 50 µl of Dig-300 Buffer (20 mM HEPES pH 7.5; 300 mM NaCl; 0.5 mM Spermidine; 1x Protease inhibitor cocktail; 0.01% Digitonin). After removing the supernatant, 50 µl of pA-Tn5 adapter complex was added to the cells with gentle vortexing and incubated for 1 hr at RT. Cells were washed twice in 1 ml Dig-300 Buffer to remove unbound pA-Tn5 protein. Cells were then resuspended in 300 µl Tagmentation buffer (10 mM MgCl_2_ in Dig-300 Buffer) and incubated at 37 °C for 1 hr. To stop tagmentation and reverse crosslink DNA, 10 µl of 0.5 M EDTA, 3 µl of 10% SDS, and 2.5 µl of 20 mg/ml Proteinase K were added to each sample and incubated at 55 °C for 1hr. To extract DNA, 300 µL Phenol-chloroform-isoamyl alcohol was added to each sample and vortexed at full speed for ∼2sec. The whole mixture was then transferred to a phase-lock tub and centrifuged at 16,000 x g for 3 min at RT. 300 μl Chloroform was added to the same phase-lock tube, inverted ∼10x, and centrifuged at 16,000 x g for 3 min at RT. The aqueous layer was collected in a fresh 1.5 mL tube containing 750 µl 100% ethanol, chilled on ice, and centrifuged at 16,000 x g for 10 min at 4 °C. The pellet was washed again in 100% ethanol and air-dried. Finally, the DNA pellet was dissolved in 25 μl 1 mM Tris-HCl pH 8, 0.1 mM EDTA. The following antibodies were used for CUT&Tag: guinea pig anti-rabbit secondary antibody (Antibodies-Online.com, #ABIN101961), H3K27ac polyclonal (Abcam, t#ab4729), N-Myc (D1V2A) Monoclonal Antibody (Cell Signaling Technology, #84406) and normal Rabbit IgG (Cell Signaling Technology, #2729).

The DNA libraries were prepared by mixing 21 µl DNA with 2 µl of a universal i5 primer and 2 µL of a uniquely barcoded i7 primer. ^63^ 25 µl NEBNext HiFi 2× PCR Master mix was added to the DNA primer mix and performed PCR amplification using the following PCR cycling conditions: 72 °C for 5 min (gap filling); 98 °C for 30 s; 13 cycles of 98 °C for 10 s and 63 °C for 10 s; final extension at 72 °C for 1 min and hold at 8 °C. Post-PCR clean-up of the libraries was performed by adding 1.3× volume of Ampure XP beads (Beckman Counter) and incubated for 10 min at RT. The tube was placed on the magnetic stand and carefully withdrawn the liquid without disturbing the beads. Beads were gently washed twice in 200 µl of 80% ethanol and resuspended in 25 µl 10 mM Tris pH 8.0 by vortexing at full speed. After 5 min, the tube was placed on the magnetic stand and carefully withdrawn the liquid into a fresh tube. The quality of the libraries was determined by checking the size distribution and concentration of libraries on an Agilent 4150 TapeStation with D1000 reagents.

### CUT&Tag Data Analysis

Sample sequence alignment and peak calling were conducted using previously described specific parameters. ^62^ CUT&Tag data were aligned to the hg38 Human genome using Bowtie2 ^64^ and peaks were called using SEACR v1.3 ^65^ from each sample using IgG controls in stringent peak calling mode. Visualization was created using deeptools. ^66^ Peaks were tested for differential expression between controls and samples with DESeq2 v1.24.0. ^67^ DESeq2 Wald tests were used to determine whether fold changes were significantly different from zero. Pre-ranked gene set enrichment analysis was conducted using shrunken fold-changes and clusterProfiler v3.12.0. ^68^ Gene ontology database was used for GSEA. ^69^ The Benjamini-Hochberg method was used to adjust p-values for false discovery in both differential expression and GSEA analyses. ^70^

### Immunoblotting

To immunoblotting, cells were lysed in modified RIPA buffer (50 mM Tris pH7.5, 150 mM NaCl, 1% NP40, 0.25% sodium deoxycholate and 1 mM EDTA) with phosphatase inhibitor (Sigma) and protease inhibitors (Millipore) for 15 min in ice followed by centrifugation. Protein concentrations were determined with the Biorad Protein Assay (Biorad). Proteins were separated on precast Novex 10% Tris-Glycine gel / NuPAGE 4-12% Bis-Tris gel at 100V using the Mini Gel Tank (Invitrogen) and were blotted onto PVDF membrane at 20V for 90 min. The following antibodies were used: N-Myc (D1V2A) Rabbit Monoclonal Antibody (Cell Signaling Technology, #84406) eIF4G1 (Cell signaling Technology, #2858), Cleaved Caspase-3 (Asp175) Antibody (Cell signaling Technology, #9661), Cleaved PARP (Asp214) (D64E10) Antibody (Cell signaling Technology, #5625), and β-Actin (13E5) Rabbit mAb HRP Conjugate (Cell signaling Technology, #5125S)

### Publicly available datasets

Data from the following publically available datasets were processed: GSE101790 (ATAC-seq in ME-1 cells), GSE101789 (ChIP-seq in ME-1 cells), GSE101788 (RNA-seq in ME-1 cells), and GSE108316 (AML patient DNase-seq),

### Statistical Analysis

For mouse leukemia survival analysis, the leukemia latency and p-values were estimated using GraphPad Prism (version 8.2.1). The p-value between groups was calculated using the log-rank test. For calculating the p-value between AML subtypes (Leukemia Gene Atlas), p-values were adjusted with Benjamini-Hochberg’s false discovery rate correction. For the rest of the analysis, the p-value was calculated using a two-tailed t-test.

## Acknowledgments

We thank Scot A. Wolfe for help with genome editing using CRISPR/CAS9 RNP approach and Benedetta Bonacci for assistance with flow cytometry.

## Authorship

Contribution: J.A.P. and P.S.P. were responsible for conceptualization; J.A.P., P.S.P., and S.S. were responsible for investigations; S.Z. and R.B. conducted bioinformatics analysis; N.Z. and S.R. provided comments on the manuscript; C.M. provided resources; J.A.P. was responsible for project administration and acquisition of funding; and J.A.P. wrote the original draft.

## References

1. Okuda, T., van Deursen, J., Hiebert, S. W., Grosveld, G. & Downing, J. R. AML1, the target of multiple chromosomal translocations in human leukemia, is essential for normal fetal liver hematopoiesis. Cell 84, 321–330, doi:10.1016/s0092-8674(00)80986-1 (1996).

2. Wang, Q. et al. The CBFbeta subunit is essential for CBFalpha2 (AML1) function in vivo. Cell 87, 697–708, doi:10.1016/s0092-8674(00)81389-6 (1996).

3. Cao, W., Adya, N., Britos-Bray, M., Liu, P. P. & Friedman, A. D. The core binding factor (CBF) alpha interaction domain and the smooth muscle myosin heavy chain (SMMHC) segment of CBFbeta-SMMHC are both required to slow cell proliferation. The Journal of biological chemistry 273, 31534–31540, doi:10.1074/jbc.273.47.31534 (1998).

4. Lukasik, S. M. et al. Altered affinity of CBF beta-SMMHC for Runx1 explains its role in leukemogenesis. Nature structural biology 9, 674–679, doi:10.1038/nsb831 (2002).

5. Castilla, L. H. et al. Failure of embryonic hematopoiesis and lethal hemorrhages in mouse embryos heterozygous for a knocked-in leukemia gene CBFB-MYH11. Cell 87, 687–696, doi:10.1016/s0092-8674(00)81388-4 (1996).

6. Castilla, L. H. et al. The fusion gene Cbfb-MYH11 blocks myeloid differentiation and predisposes mice to acute myelomonocytic leukaemia. Nature genetics 23, 144–146, doi:10.1038/13776 (1999).

7. Kuo, Y. H. et al. Cbf beta-SMMHC induces distinct abnormal myeloid progenitors able to develop acute myeloid leukemia. Cancer cell 9, 57–68, doi:10.1016/j.ccr.2005.12.014 (2006).

8. Kamikubo, Y. et al. Accelerated leukemogenesis by truncated CBF beta-SMMHC defective in high-affinity binding with RUNX1. Cancer cell 17, 455–468, doi:10.1016/j.ccr.2010.03.022 (2010).

9. Xue, L., Pulikkan, J. A., Valk, P. J. & Castilla, L. H. NrasG12D oncoprotein inhibits apoptosis of preleukemic cells expressing Cbfbeta-SMMHC via activation of MEK/ERK axis. Blood 124, 426–436, doi:10.1182/blood-2013-12-541730 (2014).

10. Zhao, L. et al. KIT with D816 mutations cooperates with CBFB-MYH11 for leukemogenesis in mice. Blood 119, 1511–1521, doi:10.1182/blood-2011-02-338210 (2012).

11. Kim, H. G. et al. FLT3-ITD cooperates with inv(16) to promote progression to acute myeloid leukemia. Blood 111, 1567–1574, doi:10.1182/blood-2006-06-030312 (2008).

12. Illendula, A. et al. Chemical biology. A small-molecule inhibitor of the aberrant transcription factor CBFbeta-SMMHC delays leukemia in mice. Science 347, 779–784, doi:10.1126/science.aaa0314 (2015).

13. Surapally, S., Tenen, D. G. & Pulikkan, J. A. Emerging therapies for inv(16) AML. Blood 137, 2579–2584, doi:10.1182/blood.2020009933 (2021).

14. Pulikkan, J. A. et al. CBFbeta-SMMHC Inhibition Triggers Apoptosis by Disrupting MYC Chromatin Dynamics in Acute Myeloid Leukemia. Cell 174, 172–186 e121, doi:10.1016/j.cell.2018.05.048 (2018).

15. DePinho, R. et al. Myc family of cellular oncogenes. Journal of cellular biochemistry 33, 257–266, doi:10.1002/jcb.240330404 (1987).

16. Delmore, J. E. et al. BET bromodomain inhibition as a therapeutic strategy to target c-Myc. Cell 146, 904–917, doi:10.1016/j.cell.2011.08.017 (2011).

17. Puissant, A. et al. Targeting MYCN in neuroblastoma by BET bromodomain inhibition. Cancer discovery 3, 308–323, doi:10.1158/2159-8290.CD-12-0418 (2013).

18. Krevvata, M. et al. Cytokines increase engraftment of human acute myeloid leukemia cells in immunocompromised mice but not engraftment of human myelodysplastic syndrome cells. Haematologica 103, 959–971, doi:10.3324/haematol.2017.183202 (2018).

19. Wunderlich, M. et al. AML xenograft efficiency is significantly improved in NOD/SCID-IL2RG mice constitutively expressing human SCF, GM-CSF and IL-3. Leukemia 24, 1785-1788, doi:10.1038/leu.2010.158 (2010).

20. Deng, W. et al. Controlling long-range genomic interactions at a native locus by targeted tethering of a looping factor. Cell 149, 1233–1244, doi:10.1016/j.cell.2012.03.051 (2012).

21. Sin-Chan, P. et al. A C19MC-LIN28A-MYCN Oncogenic Circuit Driven by Hijacked Super-enhancers Is a Distinct Therapeutic Vulnerability in ETMRs: A Lethal Brain Tumor. Cancer cell 36, 51–67 e57, doi:10.1016/j.ccell.2019.06.002 (2019).

22. Assi, S. A. et al. Subtype-specific regulatory network rewiring in acute myeloid leukemia. Nature genetics 51, 151–162, doi:10.1038/s41588-018-0270-1 (2019).

23. Xu, J. et al. Subtype-specific 3D genome alteration in acute myeloid leukaemia. Nature 611, 387–398, doi:10.1038/s41586-022-05365-x (2022).

24. Westermann, F. et al. Distinct transcriptional MYCN/c-MYC activities are associated with spontaneous regression or malignant progression in neuroblastomas. Genome biology 9, R150, doi:10.1186/gb-2008-9-10-r150 (2008).

25. Breit, S. & Schwab, M. Suppression of MYC by high expression of NMYC in human neuroblastoma cells. Journal of neuroscience research 24, 21–28, doi:10.1002/jnr.490240105 (1989).

26. Laurenti, E. et al. Hematopoietic stem cell function and survival depend on c-Myc and N-Myc activity. Cell stem cell 3, 611–624, doi:10.1016/j.stem.2008.09.005 (2008).

27. Charron, J. et al. Embryonic lethality in mice homozygous for a targeted disruption of the N-myc gene. Genes & development 6, 2248–2257, doi:10.1101/gad.6.12a.2248 (1992).

28. Ye, M. et al. C/EBPa controls acquisition and maintenance of adult haematopoietic stem cell quiescence. Nature cell biology 15, 385–394, doi:10.1038/ncb2698 (2013).

29. Wilson, A. et al. c-Myc controls the balance between hematopoietic stem cell self-renewal and differentiation. Genes & development 18, 2747–2763, doi:10.1101/gad.313104 (2004).

30. Baena, E., Ortiz, M., Martinez, A. C. & de Alboran, I. M. c-Myc is essential for hematopoietic stem cell differentiation and regulates Lin(-)Sca-1(+)c-Kit(-) cell generation through p21. Experimental hematology 35, 1333–1343, doi:10.1016/j.exphem.2007.05.015 (2007).

31. Shi, J. et al. Role of SWI/SNF in acute leukemia maintenance and enhancer-mediated Myc regulation. Genes & development 27, 2648–2662, doi:10.1101/gad.232710.113 (2013).

32. Delgado, M. D. & Leon, J. Myc roles in hematopoiesis and leukemia. Genes & cancer 1, 605–616, doi:10.1177/1947601910377495 (2010).

33. Rickman, D. S., Schulte, J. H. & Eilers, M. The Expanding World of N-MYC-Driven Tumors. Cancer discovery 8, 150–163, doi:10.1158/2159-8290.CD-17-0273 (2018).

34. Hirvonen, H., Hukkanen, V., Salmi, T. T., Pelliniemi, T. T. & Alitalo, R. L-myc and N-myc in hematopoietic malignancies. Leukemia & lymphoma 11, 197–205, doi:10.3109/10428199309086996 (1993).

35. Kawagoe, H. & Grosveld, G. C. Conditional MN1-TEL knock-in mice develop acute myeloid leukemia in conjunction with overexpression of HOXA9. Blood 106, 4269–4277, doi:10.1182/blood-2005-04-1679 (2005).

36. Fukuda, Y. et al. Upregulated heme biosynthesis, an exploitable vulnerability in MYCN-driven leukemogenesis. JCI insight 2, doi:10.1172/jci.insight.92409 (2017).

37. Liu, L. et al. MYCN contributes to the malignant characteristics of erythroleukemia through EZH2-mediated epigenetic repression of p21. Cell death & disease 8, e3126, doi:10.1038/cddis.2017.526 (2017).

38. Mundo, L. et al. Molecular switch from MYC to MYCN expression in MYC protein negative Burkitt lymphoma cases. Blood cancer journal 9, 91, doi:10.1038/s41408-019-0252-2 (2019).

39. Kawagoe, H., Kandilci, A., Kranenburg, T. A. & Grosveld, G. C. Overexpression of N-Myc rapidly causes acute myeloid leukemia in mice. Cancer research 67, 10677–10685, doi:10.1158/0008-5472.CAN-07-1118 (2007).

40. Vanden Bempt, M. et al. Aberrant MYCN expression drives oncogenic hijacking of EZH2 as a transcriptional activator in peripheral T-cell lymphoma. Blood 140, 2463–2476, doi:10.1182/blood.2022016428 (2022).

41. King, B. et al. The ubiquitin ligase Huwe1 regulates the maintenance and lymphoid commitment of hematopoietic stem cells. Nature immunology 17, 1312–1321, doi:10.1038/ni.3559 (2016).

42. Le Grand, M. et al. Interplay between MycN and c-Myc regulates radioresistance and cancer stem cell phenotype in neuroblastoma upon glutamine deprivation. Theranostics 10, 6411–6429, doi:10.7150/thno.42602 (2020).

43. Malynn, B. A. et al. N-myc can functionally replace c-myc in murine development, cellular growth, and differentiation. Genes & development 14, 1390–1399 (2000).

44. Vo, B. T. et al. The Interaction of Myc with Miz1 Defines Medulloblastoma Subgroup Identity. Cancer cell 29, 5–16, doi:10.1016/j.ccell.2015.12.003 (2016).

45. Bhat, M. et al. Targeting the translation machinery in cancer. Nature reviews. Drug discovery 14, 261–278, doi:10.1038/nrd4505 (2015).

46. Malka-Mahieu, H., Newman, M., Desaubry, L., Robert, C. & Vagner, S. Molecular Pathways: The eIF4F Translation Initiation Complex-New Opportunities for Cancer Treatment. Clinical cancer research: an official journal of the American Association for Cancer Research 23, 21–25, doi:10.1158/1078-0432.CCR-14-2362 (2017).

47. Tamburini, J. et al. Protein synthesis is resistant to rapamycin and constitutes a promising therapeutic target in acute myeloid leukemia. Blood 114, 1618–1627, doi:10.1182/blood-2008-10-184515 (2009).

48. Carroll, M. Taking aim at protein translation in AML. Blood 114, 1458–1459, doi:10.1182/blood-2009-06-224220 (2009).

49. Topisirovic, I. et al. Aberrant eukaryotic translation initiation factor 4E-dependent mRNA transport impedes hematopoietic differentiation and contributes to leukemogenesis. Molecular and cellular biology 23, 8992–9002, doi:10.1128/MCB.23.24.8992-9002.2003 (2003).

50. Nishida, Y. et al. Inhibition of translation initiation factor eIF4a inactivates heat shock factor 1 (HSF1) and exerts anti-leukemia activity in AML. Leukemia 35, 2469–2481, doi:10.1038/s41375-021-01308-z (2021).

51. Silvera, D. et al. Essential role for eIF4GI overexpression in the pathogenesis of inflammatory breast cancer. Nature cell biology 11, 903–908, doi:10.1038/ncb1900 (2009).

52. Jaiswal, P. K., Koul, S., Shanmugam, P. S. T. & Koul, H. K. Eukaryotic Translation Initiation Factor 4 Gamma 1 (eIF4G1) is upregulated during Prostate cancer progression and modulates cell growth and metastasis. Scientific reports 8, 7459, doi:10.1038/s41598-018-25798-7 (2018).

53. Comtesse, N. et al. Frequent overexpression of the genes FXR1, CLAPM1 and EIF4G located on amplicon 3q26-27 in squamous cell carcinoma of the lung. International journal of cancer 120, 2538-2544, doi:10.1002/ijc.22585 (2007).

54. Wang, H., Wang, X., Xu, L., Zhang, J. & Cao, H. Prognostic significance of MYCN related genes in pediatric neuroblastoma: a study based on TARGET and GEO datasets. BMC pediatrics 20, 314, doi:10.1186/s12887-020-02219-1 (2020).

55. Das, S. & Das, B. eIF4G-an integrator of mRNA metabolism? FEMS yeast research 16, doi:10.1093/femsyr/fow087 (2016).

56. Yamauchi, T. et al. Genome-wide CRISPR-Cas9 Screen Identifies Leukemia-Specific Dependence on a Pre-mRNA Metabolic Pathway Regulated by DCPS. Cancer cell 33, 386–400 e385, doi:10.1016/j.ccell.2018.01.012 (2018).

57. Yoshimi, A. & Abdel-Wahab, O. Targeting mRNA Decapping in AML. Cancer cell 33, 339–341, doi:10.1016/j.ccell.2018.02.015 (2018).

58. Bullinger, L. et al. Gene-expression profiling identifies distinct subclasses of core binding factor acute myeloid leukemia. Blood 110, 1291–1300, doi:10.1182/blood-2006-10-049783 (2007).

59. Shorstova, T., Foulkes, W. D. & Witcher, M. Achieving clinical success with BET inhibitors as anti-cancer agents. British journal of cancer 124, 1478–1490, doi:10.1038/s41416-021-01321-0 (2021).

60. Feng, Y. et al. SBI-0640756 Attenuates the Growth of Clinically Unresponsive Melanomas by Disrupting the eIF4F Translation Initiation Complex. Cancer research 75, 5211–5218, doi:10.1158/0008-5472.CAN-15-0885 (2015).

61. Herzog, L. O. et al. Targeting eIF4F translation initiation complex with SBI-756 sensitises B lymphoma cells to venetoclax. British journal of cancer 124, 1098–1109, doi:10.1038/s41416-020-01205-9 (2021).

62. Kaya-Okur, H. S. et al. CUT&Tag for efficient epigenomic profiling of small samples and single cells. Nature communications 10, 1930, doi:10.1038/s41467-019-09982-5 (2019).

63. Buenrostro, J. D. et al. Single-cell chromatin accessibility reveals principles of regulatory variation. Nature 523, 486–490, doi:10.1038/nature14590 (2015).

64. Langmead, B. & Salzberg, S. L. Fast gapped-read alignment with Bowtie 2. Nature methods 9, 357–359, doi:10.1038/nmeth.1923 (2012).

65. Meers, M. P., Tenenbaum, D. & Henikoff, S. Peak calling by Sparse Enrichment Analysis for CUT&RUN chromatin profiling. Epigenetics & chromatin 12, 42, doi:10.1186/s13072-019-0287-4 (2019).

66. Ramirez, F. et al. deepTools2: a next generation web server for deep-sequencing data analysis. Nucleic acids research 44, W160–165, doi:10.1093/nar/gkw257 (2016).

67. Love, M. I., Huber, W. & Anders, S. Moderated estimation of fold change and dispersion for RNA-seq data with DESeq2. Genome biology 15, 550, doi:10.1186/s13059-014-0550-8 (2014).

68. Yu, G., Wang, L. G., Han, Y. & He, Q. Y. clusterProfiler: an R package for comparing biological themes among gene clusters. Omics: a journal of integrative biology 16, 284–287, doi:10.1089/omi.2011.0118 (2012).

69. Ashburner, M. et al. Gene ontology: tool for the unification of biology. The Gene Ontology Consortium. Nature genetics 25, 25–29, doi:10.1038/75556 (2000).

70. Benjamini, Y. H., Y. Controlling the false discovery rate: a practical and powerful appoach to multiple testing. Journal of the Royal Statistical Society B, 289–300 (1995).

