## Supplementary material for "N-MYC regulates Cell Survival via eIF4G1 in inv(16) Acute Myeloid Leukemia": Figure legends, figures, table S1

**Supplementary Figures legends**

**Figure S1. MYCN is downregulated with AI-10-49 and JQ1 in inv(16) AML cells.**
(**A**). MYC and MYCN transcript levels in DMSO / AI-10-49 treated (1µM, 6 hrs) ME-1 cells by Real Time RT-PCR. Histogram representative of triplicate experiments. (**B**). N-MYC protein levels in AI-10-49 treated (6 hrs) ME-1 cells by western blot. (**C**). N-MYC transcript levels in DMSO / JQ1 treated (6 hrs) ME-1 cells by Real Time RT-PCR. Histogram representative of triplicate experiments. Error bars represent the SD. Significance was calculated using an unpaired t-test. $*p < 0.05$ or $**p < 0.005$.

**Figure S2. MYCN deletion does not lead to granulocyte differentiation in inv(16) cells.**
(**A**). The discordance plot showing the level of alignment per base between the controls (orange) and edited sample (green) in the inference window (the region around the cut site). The dotted lines represent the cut sites for gRNAs used for the experiment. (**B**). The Indel plot displaying the distribution of indels and the size of insertion or deletion (+ or -1 or more nucleotides) in the entire edited population after *MYCN* editing. (**C-F**). ME-1 cells were transfected with Cas9 and control gRNA/ pool of 3 *MYCN* gRNAs and analyzed granulocytic differentiation 11 days later by flow cytometry (C, D) and Real Time RT-PCR (E, F). Histogram representative of triplicate experiments.

**Figure S3. N-MYC silencing effectively reduces AML burden in the NSGS xenograft model.**
(**A**). Flow cytometric quantification of hCD45+ hCD33+ cells in bone marrow aspirates in NSGS mice transplanted with control/ *MYCN* edited primary inv(16) AML cells five days after transplantation. Each symbol represents a mouse. The average value of each group is shown in red. (**B**). Representative flow cytometry plots showing gating and frequency of hCD45+ hCD33+ cells in the bone marrow of NSGS mice transplanted with control/ *MYCN* edited primary inv(16) AML cells five days after transplantation.

**Figure S4. *RDME* acts as an enhancer in select AML subtypes.**
Representative examples of Integrative Genome Viewer (IGV) tracks of DNase I-seq analysis in purified healthy peripheral blood CD34+ cells and AML CD34+ cells of the indicated subtypes.

**Figure S5. CRISPR/Cas9-mediated deletion of *RDME* (A-C) and *MYCN-e2* (D-F) in ME-1 cells.**

(**A, D**). Analysis of editing efficiency by Inference of CRISPR editing (ICE). (**B, E**). The Indel plot displaying the distribution of indels and the size of insertion or deletion (+ or -1 or more nucleotides) in the entire edited population after *MYCN* enhancer editing. (**C, F**). The discordance plot showing the level of alignment per base between the control (orange) and edited sample (green) in the inference window (the region around the cut site). The dotted line represents the cut sites for upstream gRNA used for the experiment.

**Figure S6. eIF4G1 is downregulated with AI-10-49 in inv(16) AML cells.**

(**A**). EIF4G1 transcript levels in DMSO / AI-10-49 treated (9 hrs) ME-1 cells by Real Time RT-PCR. (**B**). eIF4G1 protein levels in AI-10-49 treated (1 μM) ME-1 cells by western blot. Error bars represent the SD. Significance was calculated using an unpaired t-test. *$p < 0.05$ or **$p < 0.005$.

**Figure S7. N-MYC positively regulates MYC transcript levels in inv(16) AML cells.**

(**A**). Representative examples of Integrative Genome Viewer (IGV) tracks of N-MYC CUT&Tag-seq and DNase I-seq analysis in ME-1 cells and purified human primary AML cells. (**B**). ChIP-qPCR analysis for N-MYC binding region in *MYC* promoter and a non-binding region in *MYC* locus in ME-1 cells. Histogram representative of the triplicate experiments. (**C**). ME-1 cells were transfected with Cas9 and control gRNAs/ *MYCN* gRNAs [Synthego Gene Knockout kit V2] by RNP approach and analyzed MYC transcript levels by Real Time RT-PCR. Histogram representative of triplicate experiments. Error bars represent the SD. Significance was calculated using an unpaired t-test. *$p < 0.05$ or **$p < 0.005$.

**Figure S8. CRISPR/ Cas9 mediated deletion of EIF4G1 in inv(16) AML cells.**

(**A**). ME-1 cells were transfected with Cas9 and control gRNA/ pool of 3 *EIF4G1* gRNAs [Synthego Gene Knockout kit V2] by RNP approach and analyzed editing efficiency by Inference of CRISPR editing (ICE). (**B**). The discordance plot showing the level of alignment per base between the control (orange) and edited sample (green) in the inference window (the region around the cut site). The dotted lines represent the cut sites for gRNAs used for the experiment. (**C**). The

Indel plot displaying the distribution of indels and the size of insertion or deletion (+ or -1 or more nucleotides) in the entire edited population after *EIF4G1* editing. (**D**). Flow cytometric quantification of hCD45+ hCD33+ cells in the bone marrow aspirates in NSGS mice transplanted with control/ *EIF4G1* edited primary inv(16) AML cells five days after transplantation. Each symbol represents a mouse. The average value of each group is shown in red. (**E**). Representative flow cytometry plots showing gating and frequency of hCD45+ hCD33+ cells in the bone marrow of NSGS mice transplanted with control/ *EIF4G1* edited primary inv(16) AML cells five days after transplantation.

Figure S1

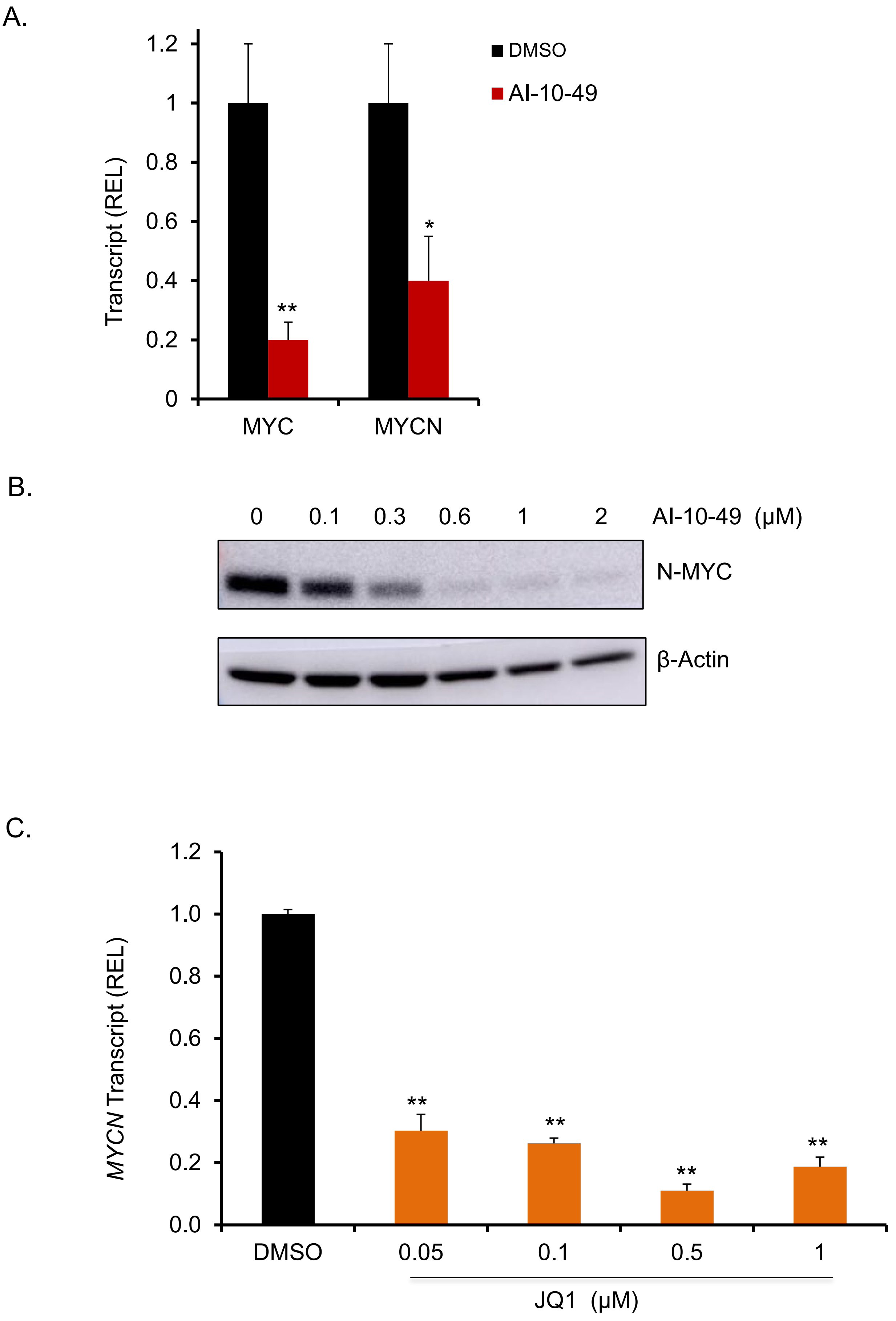
A.
1.2
1
0.8
0.6
0.4
0.2
0
Transcript (REL)
DMSO
AI-10-49
**
*
MYC
MYCN
B.
0
0.1
0.3
0.6
1
2
AI-10-49 (μM)
N-MYC
β-Actin
C.
1.2
1.0
0.8
0.6
0.4
0.2
0.0
MYCN Transcript (REL)
**
**
**
**
DMSO
0.05
0.1
0.5
1
JQ1 (μM)

Figure S2

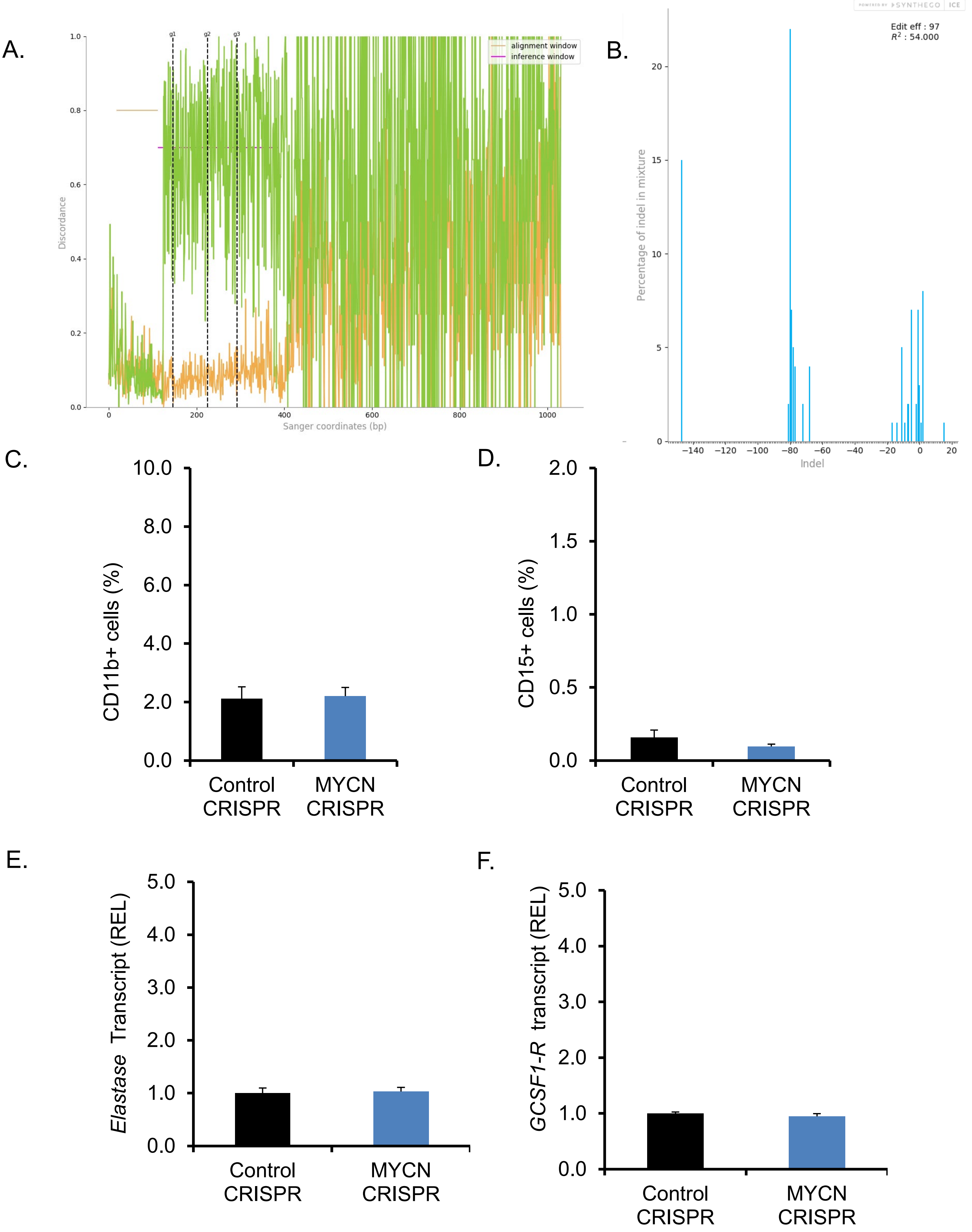

A.
g1
g2
g3
alignment window
inference window
Discordance
Sanger coordinates (bp)
B.
SYNTHEGO ICE
Edit eff : 97
$R^2$ : 54.000
Percentage of indel in mixture
Indel
C.
CD11b+ cells (%)
Control CRISPR
MYCN CRISPR
D.
CD15+ cells (%)
Control CRISPR
MYCN CRISPR
E.
*Elastase* Transcript (REL)
Control CRISPR
MYCN CRISPR
F.
*GCSF1-R* transcript (REL)
Control CRISPR
MYCN CRISPR

Figure S3

A.

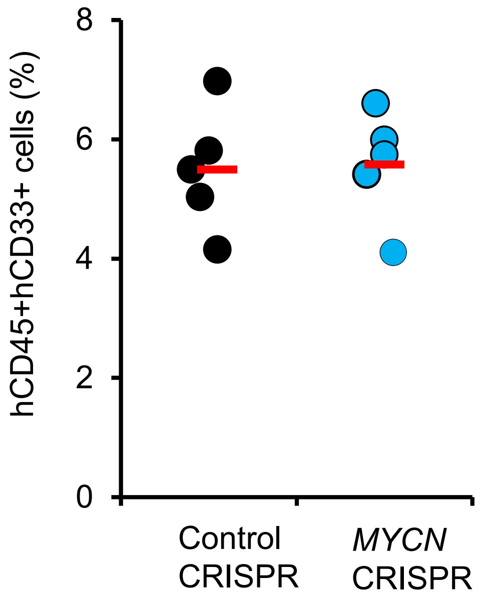

hCD45+hCD33+ cells (%)
8
6
4
2
0
Control
CRISPR
*MYCN*
CRISPR


B.

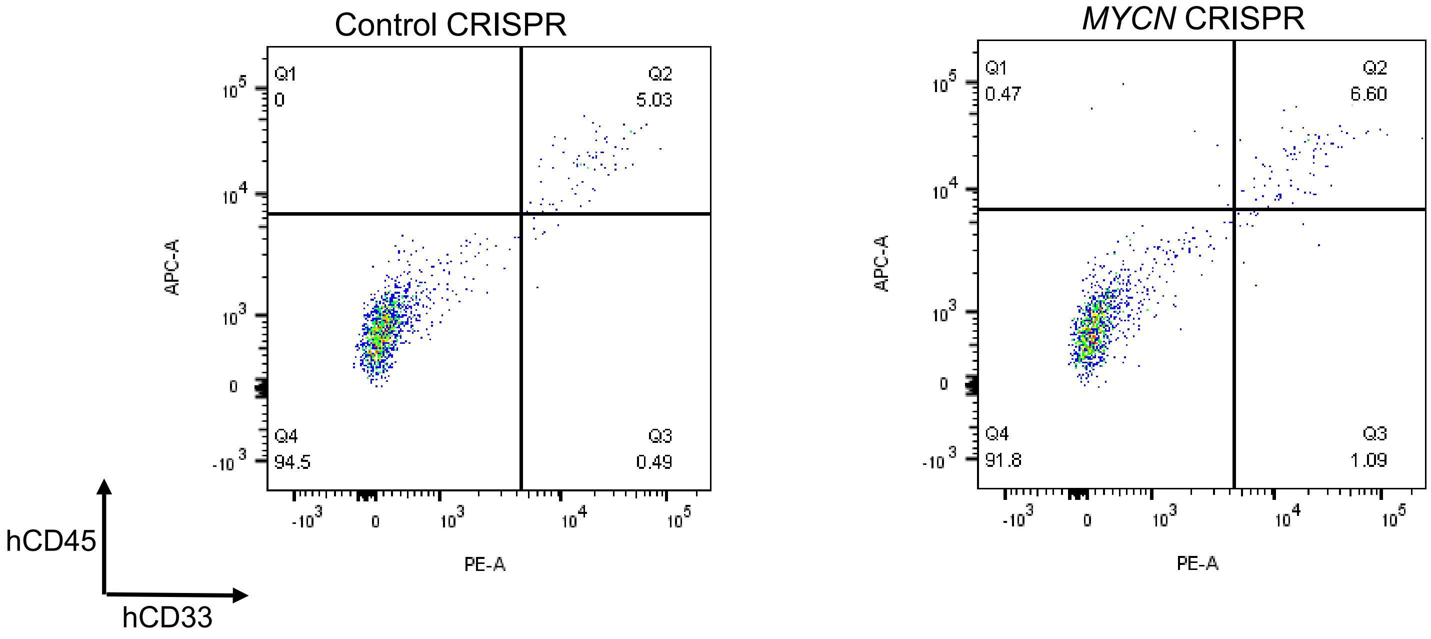

Control CRISPR
*MYCN* CRISPR
Q1
0
Q2
5.03
Q4
94.5
Q3
0.49
Q1
0.47
Q2
6.60
Q4
91.8
Q3
1.09
APC-A
PE-A
hCD45
hCD33

Figure S4

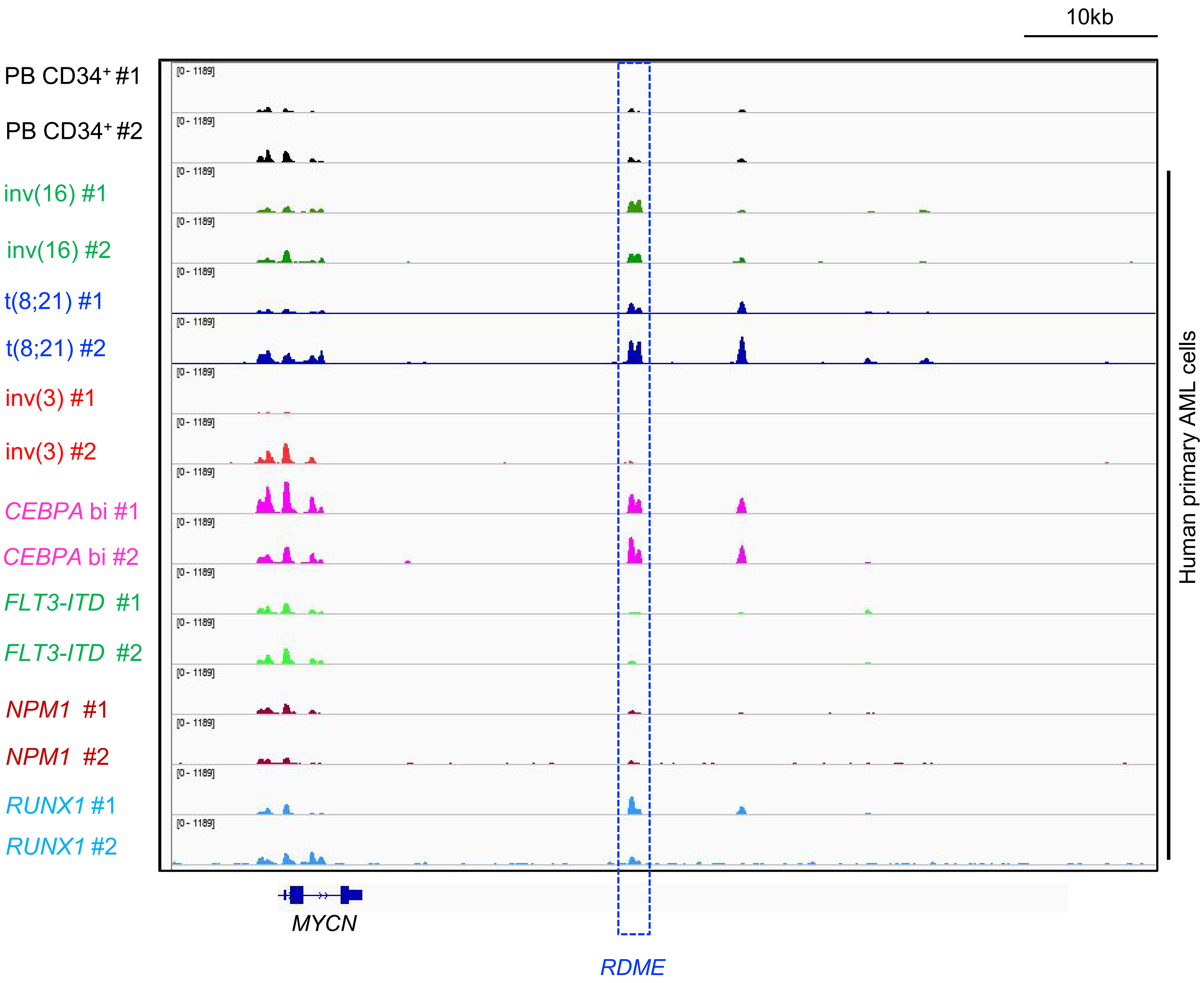

10kb
PB CD34+ #1
PB CD34+ #2
inv(16) #1
inv(16) #2
t(8;21) #1
t(8;21) #2
inv(3) #1
inv(3) #2
CEBPA bi #1
CEBPA bi #2
FLT3-ITD #1
FLT3-ITD #2
NPM1 #1
NPM1 #2
RUNX1 #1
RUNX1 #2
[0 - 1189]
Human primary AML cells
MYCN
RDME

Figure S5

A.

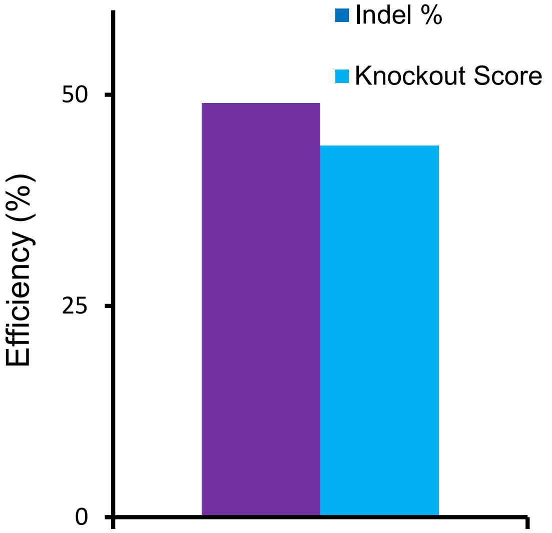


B.

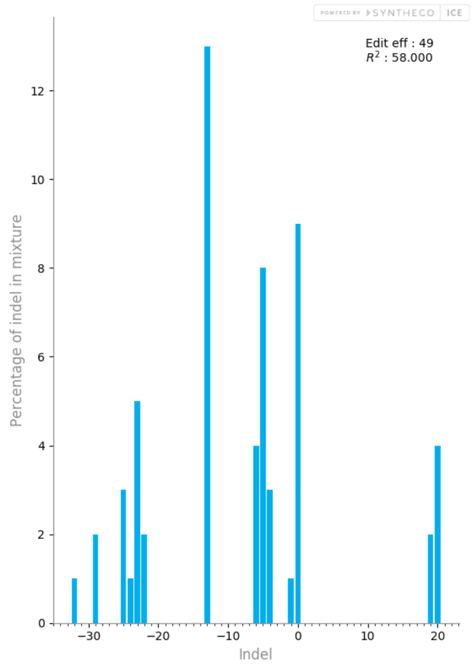


C.

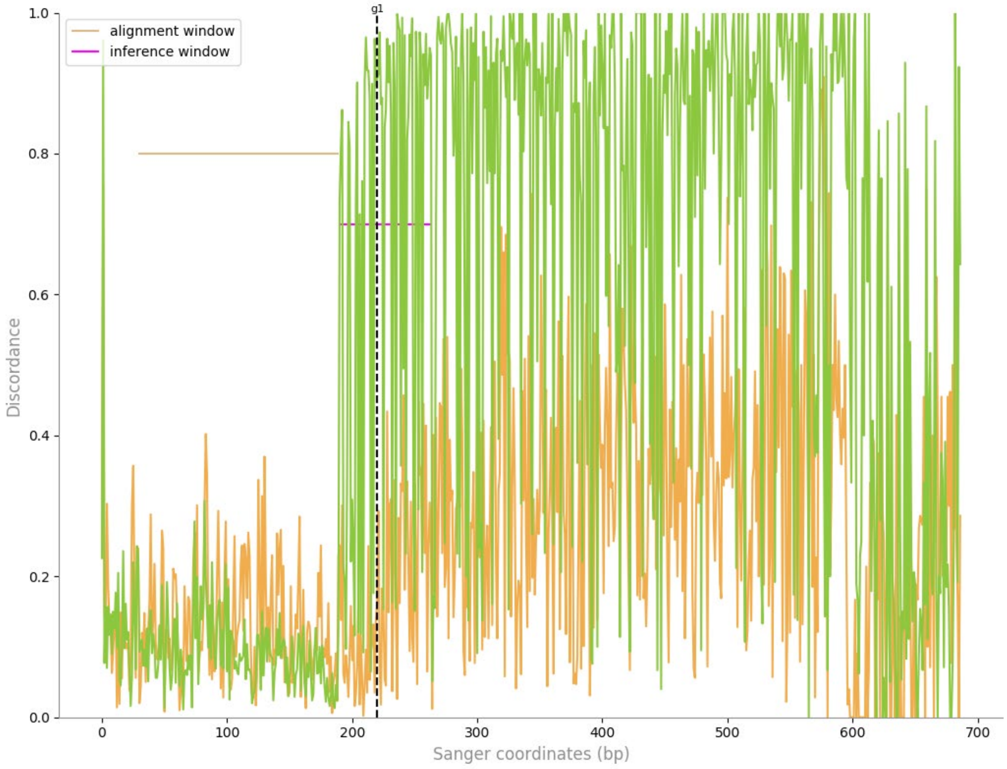

Figure S5

D.

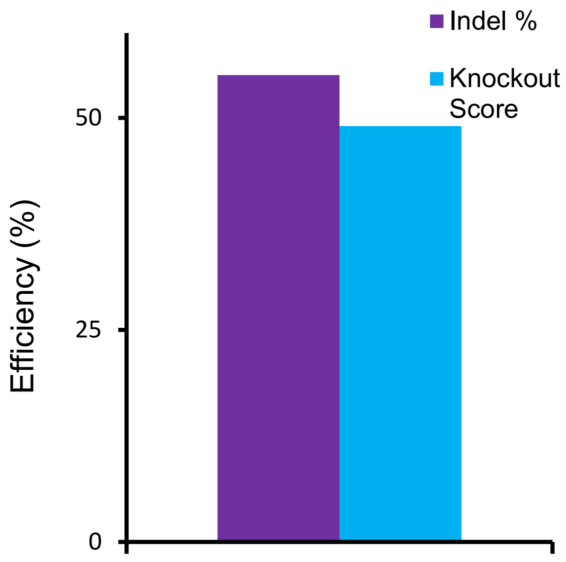


E.

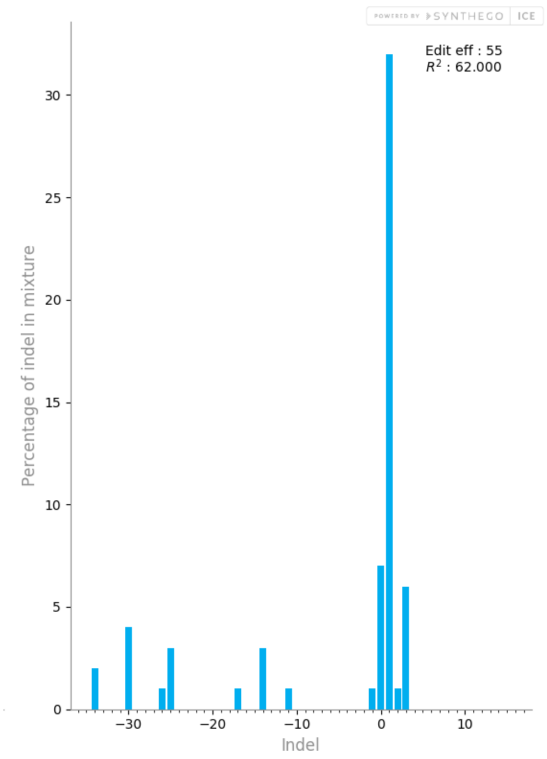


F.

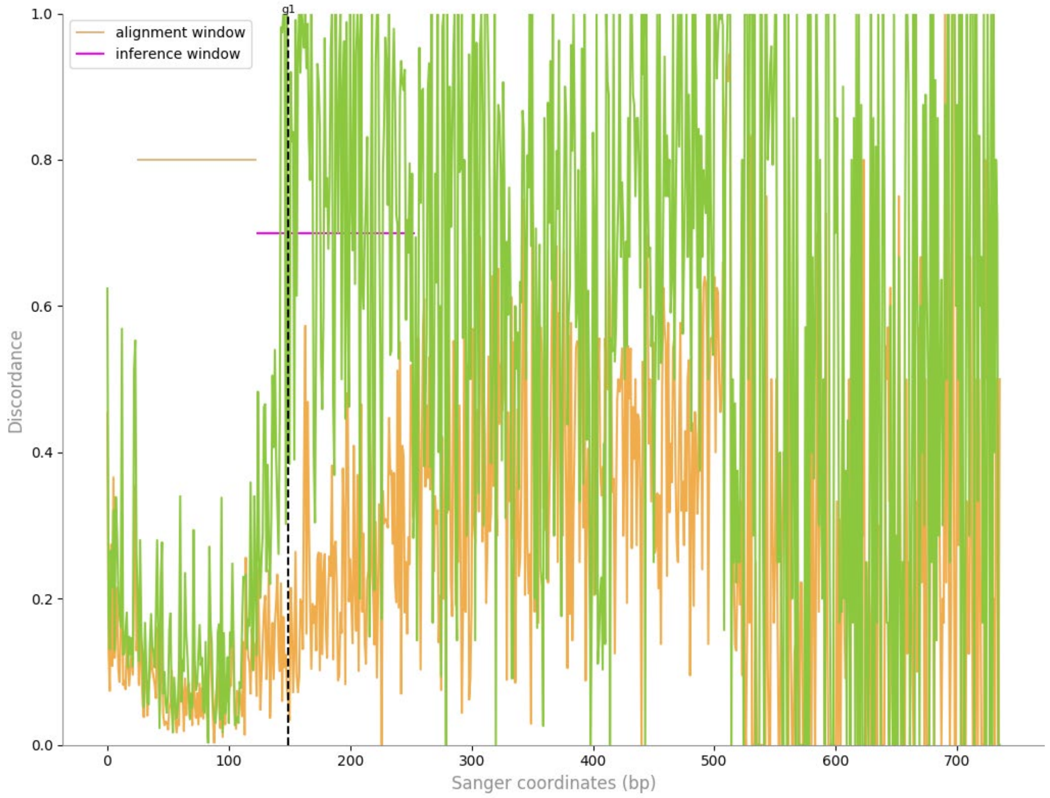

Figure S6

A.

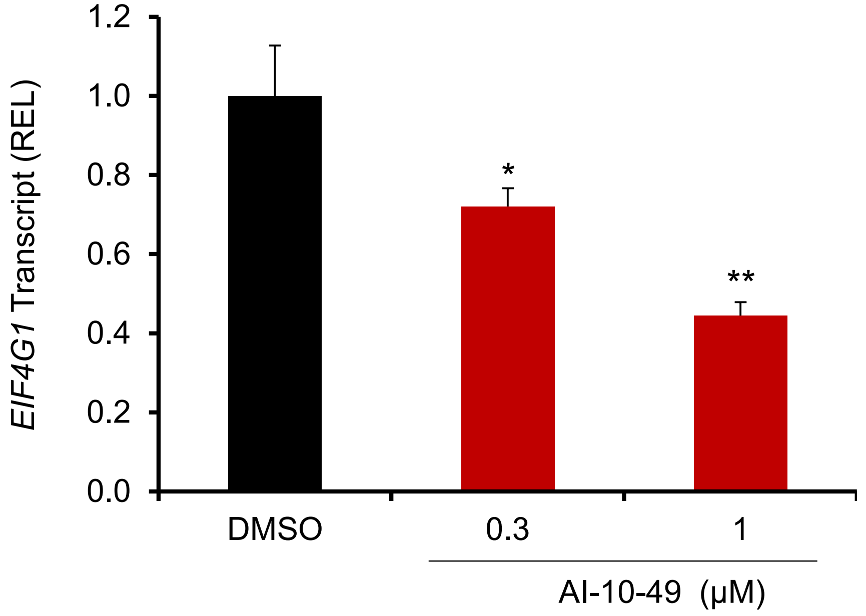

1.2
1.0
0.8
0.6
0.4
0.2
0.0
EIF4G1 Transcript (REL)
*
**
DMSO
0.3
1
AI-10-49 (µM)


B.

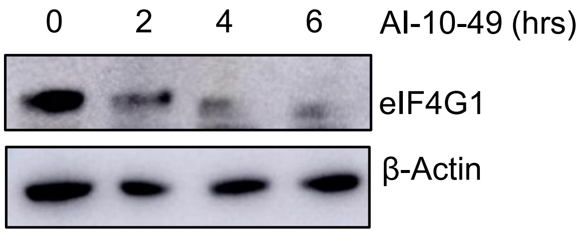

0
2
4
6
AI-10-49 (hrs)
eIF4G1
β-Actin

Figure S7

A.

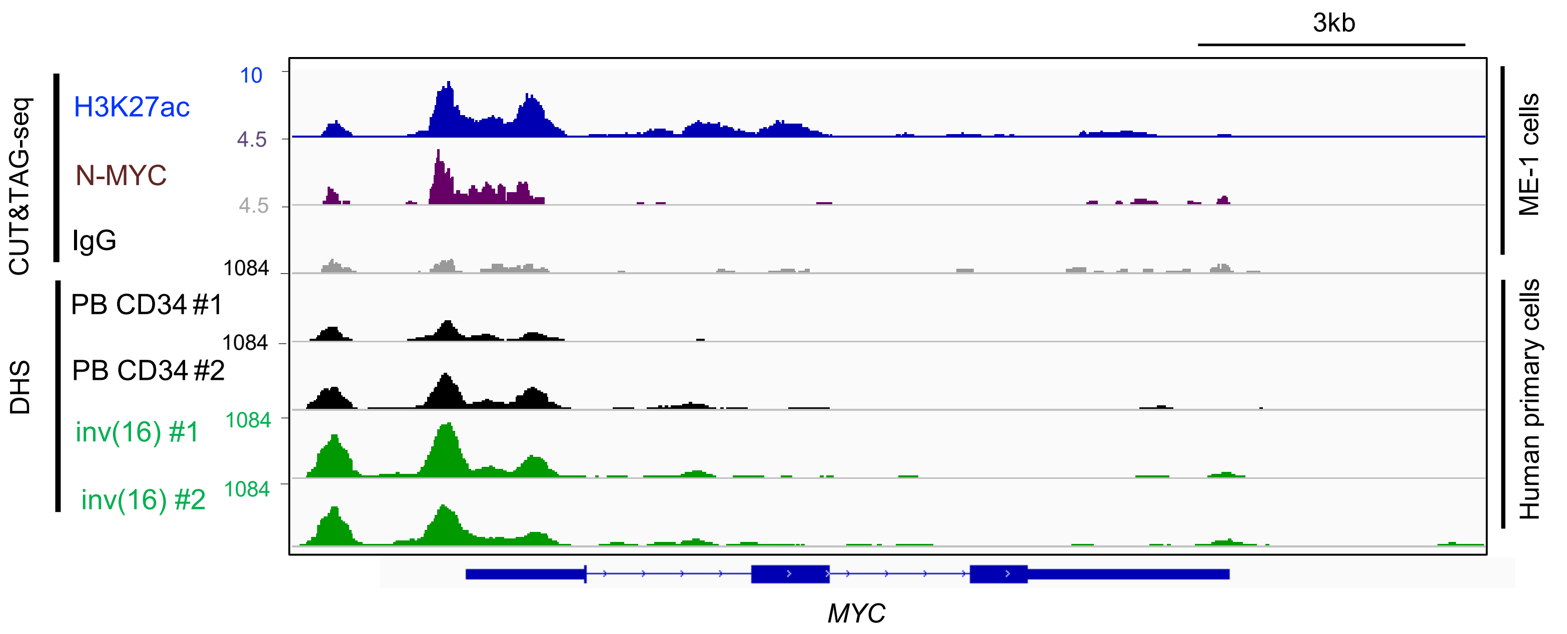


B.

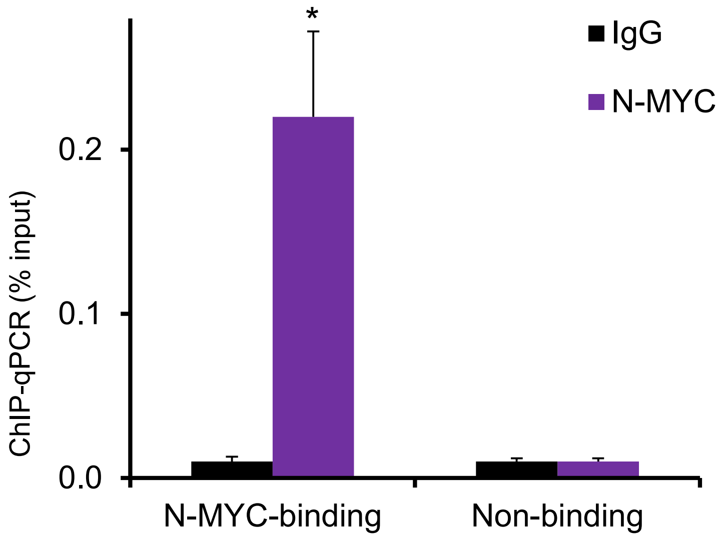


C.

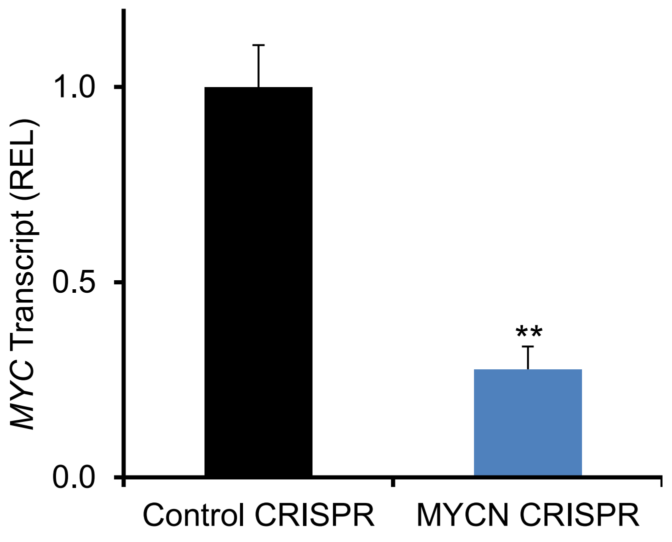

Figure S8

A.

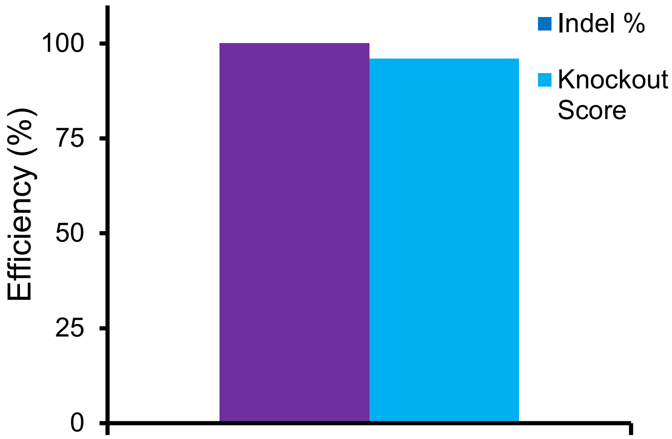


B.

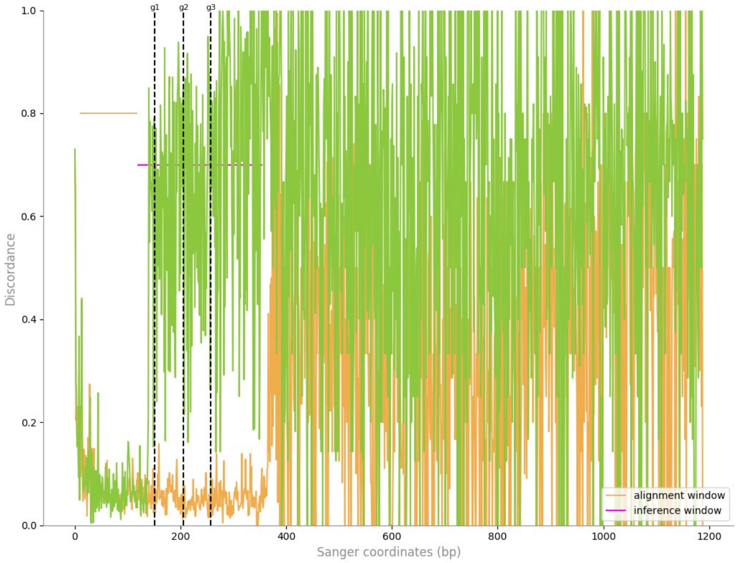


C.

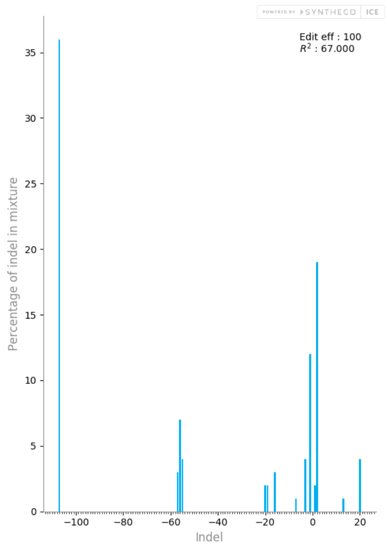


D.

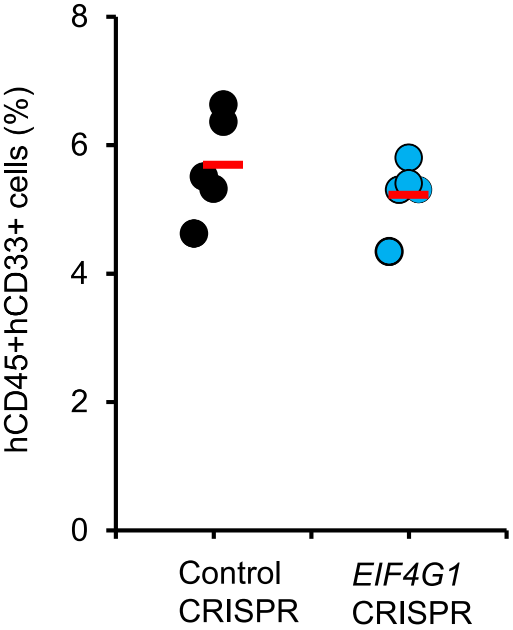

Figure S8

E.

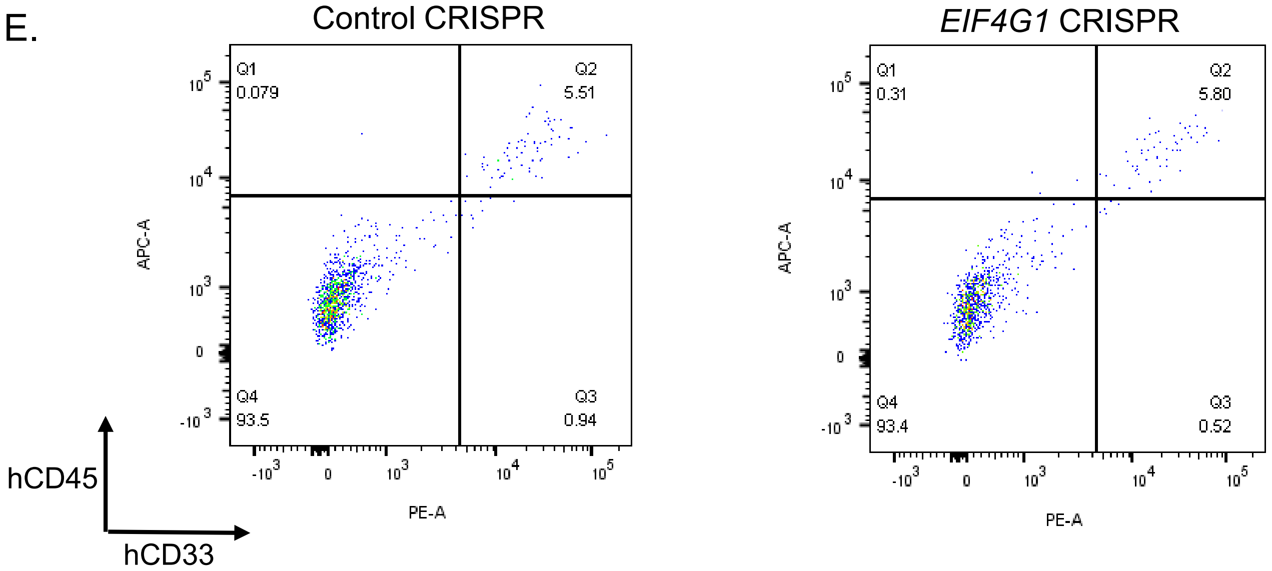
Control CRISPR
EIF4G1 CRISPR
Q1
0.079
Q2
5.51
Q4
93.5
Q3
0.94
Q1
0.31
Q2
5.80
Q4
93.4
Q3
0.52
APC-A
PE-A
hCD45
hCD33

**Table S1.** List of Oligonucleotide Sequences.

| APPLICATION | NAME | SEQUENCE (5'-3') |
|---|---|---|
| Human MYCN mRNA quantification | MYCN For | ACCACAAGGCCCTCAGTACCTC |
| Human MYCN mRNA quantification | MYCN Rev | GATGACACTCTTGAGCGGACGTGG |
| Human MYC mRNA quantification | MYC For | GCAGCTGCTTAGACGCTGGATTTT |
| Human MYC mRNA quantification | MYC Rev | GCAGCAGCTCGAATTTCTTCCAGA |
| Human EIF4G1 mRNA quantification | EIF4G1 For | TTCAGAATCCCAGCCTTCGT |
| Human EIF4G1 mRNA quantification | EIF4G1 Rev | GAGTGGGTTCTGGAGAGAGG |
| Human Elastase mRNA quantification | Elastase For | CCACCCGGCAGGTGTTC |
| Human Elastase mRNA quantification | Elstase Rev | GTGGCCGACCCGTTGAG |
| Human GCSF1-R mRNA quantification | G-CSFR For | AAGAGCCCCCTTACCCACTACACCATCTT |
| Human GCSF1-R mRNA quantification | G-CSFR Rev | TGCTGTGAGCTGGGTCTGGGACACTT |
| Human *GAPDH* mRNA quantification | GAPDH For | AGGACTCATGACCACAGT |
| Human *GAPDH* mRNA quantification | GAPDH Rev | CAGTAGAGGCAGGGATGA |
| ChIP assay for *EIF4G1* promoter (N-MYC binding) | ChIP EIF4G1 PR For | AAACACAGAGGGGAATCGGT |
| ChIP assay for *EIF4G1* promoter (N-MYC binding) | ChIP EIF4G1 PR Rev | GTAGTCCACAACCATTTCCGG |
| ChIP assay for *EIF4G1* promoter (Non-binding) | ChIP EIF4G1 NB For | GTGCTCTCGAACTCCTGACT |
| ChIP assay for *EIF4G1* promoter (Non- binding) | ChIP EIF4G1 NB Rev | CTGCACTCTAGCCTGGATGA |
| ChIP assay for *MYC* promoter (N-MYC binding) | ChIP MYC PR For | AACCTCCCTCTCGCCCTAG |
| ChIP assay for *MYC* promoter (N-MYC binding) | ChIP MYC PR Rev | CCTTCCACCCAGACTGAGTC |
| ChIP assay for *MYC* promoter (Non-binding) | ChIP MYC NB For | GATGACTCGCACTTGTATTCCC |
| ChIP assay for *MYC* promoter (Non- binding) | ChIP MYC NB Rev | CAGTCTGGTCATGTTGCCCA |
| ChIP assay for *MYCN RDME* enhancer (RUNX1 binding) | ChIP MYCN-RX1 For | TAAGGAATAGGCCCCTGGTA |
| ChIP assay for *MYCN RDM*E enhancer (RUNX1 binding) | ChIP MYCN-RX1 Rev | AGCCCTGCTGACAGCTTGA |
| ChIP assay for *MYCN-e2* enhancer (RUNX1 binding) | ChIP MYCN-Ctr For | TATGATGGGAGGAGCTGGAG |
| ChIP assay for *MYCN-e2* enhancer (RUNX1 binding) | ChIP MYCN-Ctr Rev | GGCATCACCTGGAAGTTTGT |
| CRIPSR/Cas9 *MYCN RDME* enhancer deletion | MYCN sg1 For | CACCGCACAATAAGGAATAGGCCCC |
| CRIPSR/Cas9 *MYCN RDME* enhancer deletion | MYCN sg1 Rev | AAACGGGGCCTATTCCTTATTGTGC |
| CRIPSR/Cas9 *MYCN RDME* enhancer deletion | MYCN sg2 For | CACCGTGGGCGTCTCCCGTAGCTTA |
| CRIPSR/Cas9 *MYCN RDME* enhancer deletion | MYCN sg2 Rev | AAACTAA GCT ACG GGA GAC GCC CAC |
| CRIPSR/Cas9 *MYCN-e2* enhancer deletion | MYCN sg33 For | CACCGTGTTATATCATCATTATGAT |
| CRIPSR/Cas9 *MYCN-e2* enhancer deletion | MYCN sg33 Rev | AAACATCATAATGATGATATAACAC |
| CRIPSR/Cas9 *MYCN-e2* enhancer deletion | MYCN sg34 For | CACCGGGGCCACATTGTCAATAACA |
| CRIPSR/Cas9 *MYCN-e2* enhancer deletion | MYCN sg34 Rev | AAACTGTTATTGACAATGTGGCCCC |
| ICE for analyzing *MYCN RDME* enhancer deletion | MYCN Rx1 ICE-For | AGAACCTGCCCTTCATCCC |
| ICE for analyzing *MYCN RDME* enhancer deletion | MYCN Rx1 ICE-Rev | TGACCTCCAGTACCCCAGA |
| ICE for analyzing *MYCN-e2* enhancer deletion | MYCN Ctr ICE-For | GTCTTGGCCACTTGTCAGC |
| ICE for analyzing *MYCN-e2* enhancer deletion | MYCN Ctr ICE-Rev | TTGGGTGTGCAGGGAGAA |
| CM-sgRNAs for *MYCN* deletion by CRISPR/Cas9 RNP | MYCN CM-sgRNA#1 | CUUGCAGAUCAUGCCCGGCA |
| CM-sgRNAs for *MYCN* deletion by CRISPR/Cas9 RNP | MYCN CM-sgRNA#2 | GACGAAGAUGACUUCUACUU |
| CM-sgRNAs for *MYCN* deletion by CRISPR/Cas9 RNP | MYCN CM-sgRNA#3 | CUGGGCGACAGCGGGGGCGU |
| ICE for analyzing *MYCN* deletion | MYCN-Syn-For | CTCTCCGGTGTGTCTGTCG |
| ICE for analyzing *MYCN* deletion | MYCN-Syn-Rev | CTGGAGGATGACCGGGTTG |
| CM-sgRNAs for *EIF4G1* deletion by CRISPR/Cas9 RNP | EIF4G1 CM-sgRNA#1 | AUAGUUGUCAUAGUGUCCCC |
| CM-sgRNAs for *EIF4G1* deletion by CRISPR/Cas9 RNP | EIF4G1 CM-sgRNA#2 | AUAUGGCUCCCCAGUUUCAC |
| CM-sgRNAs for *EIF4G1* deletion by CRISPR/Cas9 RNP | EIF4G1 CM-sgRNA#3 | UUCUACUUCCAGUAUGGGUU |
| ICE for analyzing *EIF4G1* deletion | EIF4G1-Syn-For | CCTATGGTGCAGATGACCGG |
| ICE for analyzing *EIF4G1* deletion | EIF4G1-Syn-Rev | GCTTGACTCCACCTCTGGTT |
